# Genetic variation in a complex polyploid: unveiling the dynamic allelic features of sugarcane

**DOI:** 10.1101/361089

**Authors:** Danilo Augusto Sforça, Sonia Vautrin, Claudio Benicio Cardoso-Silva, Melina Cristina Mancini, María Victoria Romero da Cruz, Guilherme da Silva Pereira, Mônica Conte, Arnaud Bellec, Nair Dahmer, Joelle Fourment, Nathalie Rodde, Marie-Anne Van Sluys, Renato Vicentini, Antônio Augusto Franco Garcia, Eliana Regina Forni-Martins, Monalisa Sampaio Carneiro, Hermann Paulo Hoffmann, Luciana Rossini Pinto, Marcos Guimarães de Andrade Landell, Michel Vincentz, Helene Berges, Anete Pereira Souza

## Abstract

**Background:** Sugarcane (*Saccharum spp*.) is highly polyploid and aneuploid. Modern cultivars are derived from hybridization between *S. officinarum* and *S. spontaneum*. This combination results in a genome exhibiting variable ploidy among different loci, a huge genome size (approximately 10 Gb) and a high content of repetitive regions. Gene expression mechanisms are poorly understood in these cultivars. An approach using genomic, transcriptomic and genetic mapping can improve our knowledge of the behavior of genetics in sugarcane.

**Results:** The hypothetical *HP600* and centromere protein C (*CENP-C*) genes from sugarcane were used to elucidate the allelic expression and genomic and genetic behavior of this complex polyploid. The genomically side-by-side genes *HP600* and *CENP-C* were found in two different homeologous chromosome groups with ploidies of eight and ten. The first region (Region01) was a *Sorghum bicolor* ortholog with all haplotypes of *HP600* and *CENP-C* expressed, but *HP600* exhibited an unbalanced haplotype expression. The second region (Region02) was a scrambled sugarcane sequence formed from different noncollinear genes containing duplications of *HP600* and *CENP-C* (paralogs). This duplication occurred before the *Saccharum* genus formation and after the separation of sorghum and sugarcane, resulting in a nonexpressed *HP600* pseudogene and a recombined fusion version of *CENP-C* and orthologous gene Sobic.003G299500 with at least two chimerical gene haplotypes expressed. The genetic map construction supported the difficulty of mapping markers located in duplicated regions of complex polyploid genomes.

**Conclusion:** All these findings describe a low synteny region in sugarcane, formed by events occurring in all members of the *Saccharum* genus. Additionally, evidence of duplicated and truncate gene expression and the behavior of genetic markers in a duplicated region was found. Thus, we describe the complexity involved in sugarcane genetics and genomics and allelic dynamics, which can be useful for understanding the complex polyploid genome.

## Background

The *Saccharum* species are C4 grass and present a high level of ploidy. *S. officinarum* L. is octaploid (2n = 80), with x = 10 chromosomes, while *S. spontaneum* L. has x = 8 but presents great variation in the number of chromosomes, with main cytotypes of 2n = 62, 80, 96, 112 or 128. The modern sugarcane cultivars originated from hybridization between these two species and are considered allopolyploid hybrids [1, 2]. The development of these cultivars involved the process of ‘nobilization’ of the hybrid, with successive backcrosses using *S. officinarum* as the recurrent parent [3]. The resulting hybrids are highly polyploid and aneuploid [4-6] and have an estimated whole genome size of 10 Gb [7]. An in situ hybridization study has shown that the genomes of the commercial hybrids consist of 10-20% chromosomes from *S. spontaneum* and 5-17% recombinant chromosomes between the two species, while the remaining majority of the genome consists of chromosomes from *S. officinarum* [8, 9].

Molecular evidence suggests that polyploid genomes can present dynamic changes in DNA sequence and gene expression, probably in response to genomic shock (genomic remodeling due to the activation of previously deleted heterochromatic elements), and this phenomenon is implicated in epigenetic changes in homologous genes due to intergenomic interactions [10]. The evolutionary success of polyploid species is related to their ability to present greater phenotypic novelty than is observed in their diploid or even absent in parents [11]. Among other factors, this increase in the capacity for phenotypic variation capacity may be caused by regulation of the allelic dosage [12].

The Brazilian sugarcane variety SP80-3280 is derived from a cross between the varieties SP71-1088□×□H57-5028 and is resistant to brown rust, caused by *Puccinia melanocephala* [13]. SP80-3280, which is one of the main Brazilian cultivars [14], was chosen for transcriptome sequencing by SUCEST-FUN [15] and RNAseq [16-18]. Biparental crossing of SP80-3280 has also been used to analyze rust resistance [19], quantitative trait loci (QTL) mapping [20] and genotyping by sequencing (GBS) [21]. A Brazilian initiative [22] is producing a gene-space genome sequence from SP80-3280, and a draft sugarcane genome based on whole-genome shotgun sequencing was produced [23]. In addition, QTL gene synteny from sorghum has been used to map corresponding bacterial artificial chromosomes (BACs) in SP80-3280 [24].

Three BAC libraries for different sugarcane varieties have been constructed. The oldest one is for the French variety R570 [25] and contains 103,296 clones with an average insert size of 130 kb, representing 1.2 total genome equivalents. A mix of four individuals derived from the self-fertilization of the elite cultivar R570 (pseudo F2) was reported by Le Cunff et al. [26] and contains 110,592 clones with an average insert size of 130 kb, representing 1.4x coverage of the whole genome. In addition, a library of SP80-3280 published by Figueira et al. [27] contains 36,864 clones with an average insert size of 125 kb, representing 0.4 total genome equivalent coverage.

Sugarcane and sorghum (*Sorghum bicolor* (L.) Moench) share a high level of collinearity, gene structure and sequence conservation. De Setta et al. [28] contributed to understanding the euchromatic regions from R570 and a few repetitive-rich regions, such as centromeric and ribosomal regions, other than defining a basic transposable element dataset. The genomic similarity between sugarcane and sorghum has been frequently used to characterize the sugarcane genome [24, 29-32], demonstrating the high synteny of sugarcane × sorghum and high gene structure retention among the different sugarcane homeologs. In addition, these works contribute to understanding the genomic and evolutionary relationships among important genes in sugarcane using BAC libraries.

The segregation of chromosomes during cell division is facilitated by the attachment of mitotic spindle microtubules to the kinetochore at the chromosomal centromere. Only CenH3 (histone H3) and *CENP-C* (centromere protein C) have been shown to bind centromeric DNA [33]. The centromere is marked with the histone H3 variant CenH3 (*CENP-A* in human), and *CENP-C* forms part of the inner kinetochore. The *CENP-C* “central domain” makes close contact with the acidic patch of histones H2A/H2B, and the highly conserved “*CENP-C* motif” senses both the acidic patch and recognizes the hydrophobicity of the otherwise nonconserved CenH3 tail, supporting a conserved mechanism of centromere targeting by the kinetochore [34-36]. Sandmann et al. [36] reported that in *Arabidopsis thaliana,* KNL2, a protein with a *CENPC-k* motif, recognizes centromeric nucleosomes such as the *CENP-C* protein. The CENP-C gene genomic structure in sugarcane has not been detailed.

Genome organization and expression dynamics are poorly understood in complex polyploid organisms, such as sugarcane, mainly because reconstructing large and complex regions of the genome is a challenge. However, an intriguing question is how such a complex genome can function while handling different copy numbers of genes, different allelic dosages and different ploidies of its homo/homeolog groups. For that reason, we examined the genome, transcriptome, evolutionary pattern and genetic interactions/relationships of a *CENP-C*-containing region in a genomic region of the SP80-3280 sugarcane variety (a *Saccharum* hybrid). First, we defined the genome architecture and evolutionary relationships of two physically linked genes, *HP600* (unknown function) and *CENP-C*, in detail. Second, we used the sugarcane SP80-3280 transcriptome to investigate transcription and genomic interactions in each gene (*HP600* and *CENP-C*). Ultimately, we used SNP distribution from these genes to compare the genetic and physical maps.

## Results

### BAC library construction

The BAC library from the sugarcane variety SP80-3280 resulted in 221.184 clones, arrayed in 576 384-well microtiter plates, with a mean insert size of 110 kb. This BAC library is approximately 2.4 genome equivalents (10 Gb) and 26 monoploid genome equivalents (930 Mb, [27]). For the sugarcane variety IACSP93-3046, the library construction resulted in 165.888 clones arrayed in 432 384-well microtiter plates, with a mean insert size of 110 kb, which is approximately 1.8 genome equivalents and 19 monoploid genome equivalents.

BAC-end sequencing (BES) results in an overview of the genome and validates the clones obtained through library construction. The SP80-3280 BAC library yielded 650 (84.6%) good BES sequences, of which 319 sequences have repetitive elements, and 92 exhibited similarities with sorghum genes. Excluding hits for more than one gene (probably duplicated genes or family genes), 65 sequences could be mapped to the *S. bicolor* genome (see Supplemental Figure 1, Additional File 1). The BAC library for IACSP93-3046 yielded 723 (94%) good BES sequences, of which 368 sequences exhibited the presence of repetitive sequences, and 111 exhibited a similarity with some gene. Excluding genes with hits with more than one gene, 74 of the sequences could be mapped to the *S. bicolor* genome (see Supplemental Figure 1, Additional File 1).

### BAC annotation

The gene HP600 was used as target gene and showed strong evidence of being a single-copy gene when the transcripts of HP600 of sorghum, rice and sugarcane was compared. Twenty-two BAC clones from the SP80-3280 library that had the target gene *HP600* (NCBI from MH463467 to MH463488) and a previously sequenced BAC (Mancini et al. [24]; NCBI Accession Number MF737011) were sequenced by Roche 454 sequencing (see Supplemental Table 1, Additional File 1). The BACs varied in size from 48 kb (Shy171E23) to 162 kb (Shy432H18), with a mean size of 109 kb. The BACs were compared, and BACs with at least 99% similarity were considered the same haplotype (Figures 1 and 2), resulting in sixteen haplotypes. Indeed, the possibility of one homeolog being more than 99% similar to another exists, but a real haplotype cannot be distinguished from an assembly mismatch.

**Figure 1.**
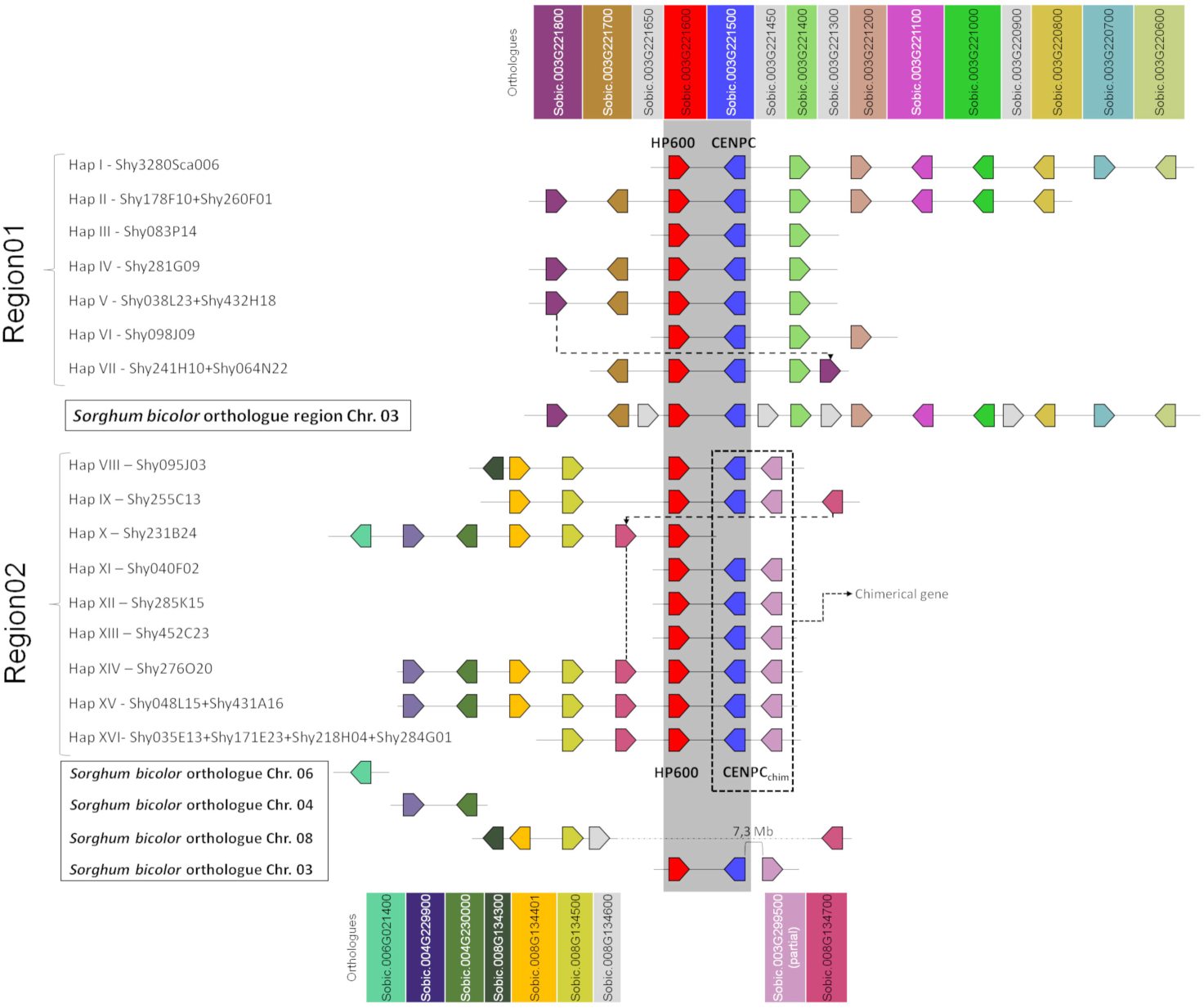
Schematic representation of the sugarcane BAC haplotypes from Region01 and Region02. Squares of the same color represent sugarcane genes orthologous to *Sorghum bicolor* genes. Dotted lines connect the homologous genes in sugarcane at different positions. In sugarcane Region02, the *CENP-C* haplotypes in Region02 are represented by two squares (blue and pink), where each square represents a partial gene fusion. The dark gray strip represents the shared region from Region01 and Region02 (duplication). The genes in light gray (from *S. bicolor*) are not found in the sugarcane BACs. The representation is not to scale. The orientation of transcription is indicated by the direction of the arrow at the end of each gene.

**Figure 2.**
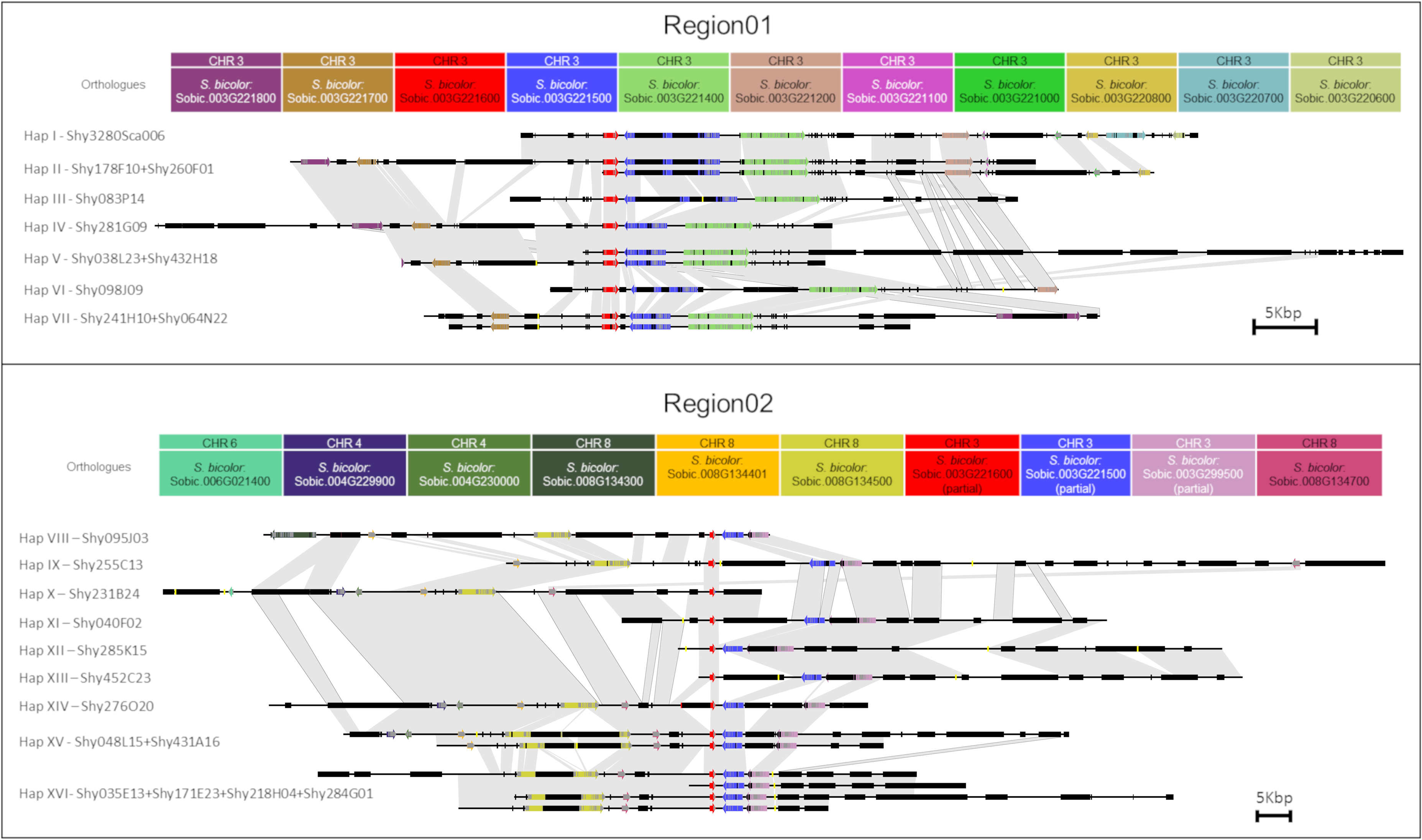
Representation of each sugarcane BAC from Region01 and Region02. Arrows and rectangles of the same color represent the homologous genes in sugarcane. Black rectangles represent repeat regions. Yellow lines represent gaps. Similar regions are represented by a gray shadow connecting the BACs. The orientation of transcription is indicated by the direction of the arrow at the end of each gene. Scale representation.

Comparisons of the BAC sequences against the sugarcane SP80-3280 genome draft using BLASTN [23] resulted in matches within gene regions, but no genome contig covered a whole BAC, and the BAC transposable elements (TEs) matched with several genome contigs (see Supplemental Figure 2, Additional File 1). The matches with genes provide further support for our assembly process.

The BACs were first annotated regarding the TEs. The TEs accounted for 21% to 65% of the sequenced bases with a mean of 40% (see Supplemental Table 1, Additional File 1). Annotation of the TEs in the 22 BACs revealed 618 TEs (220 TEs were grouped in the same type) with sizes ranging from 97 bp to 18,194 bp.

Gene annotation (see Supplemental Tables 2 and 3, Additional File 1) resulted in three to nine genes per BAC with a mean of five genes per BAC (see Supplemental Table 1, Additional File 1). The gene Sobic.003G221500, which was used to screen the library, codes for a hypothetical protein called *HP600* in sugarcane that has been found to be expressed in sorghum and rice. A phylogenetic analysis using sorghum, rice and *Arabidopsis thaliana* transcripts revealed that this gene is probably a single-copy gene. The gene Sobic.003G221600 is a *CENP-C* ortholog in sugarcane (*S. officinarum*, haplotypes CENP-C1 and CENP-C2, described by Talbert et al. [33]). The *HP600* and *CENP-C* sugarcane genes, as in *S. bicolor* and *Oryza sativa* L, were found to be side by side in the sugarcane haplotypes.

### Relationship between Region01 and Region02

Annotation of *HP600* and *CENP-C* in the sixteen BAC haplotypes revealed two groups of BACs. One group had the expected exon/intron organization when compared with *S. bicolor HP600* (five exons in sorghum) and *CENP-C* (fourteen exons in sorghum). This region was further designated as Region01 (see Supplemental Table 1, Additional File 1 - 10 BACs and 7 haplotypes – Figure 1 - haplotype I to haplotype VII). The other group was found to have fewer exons than expected (when compared with *S. bicolor*) for both *HP600* and *CENP-C*, and it was designated Region02 (see Supplemental Table 1, Additional File 1 - 13 BACs and 9 haplotypes – Figure 1 - haplotype VIII to haplotype XVI).

A comparison of the BAC haplotypes from Region01 and Region02 revealed an 8-kb shared region. The 8-kb duplication spanned from the last three exons of *HP600* to the last seven exons of *CENP-C*. *HP600* and *CENP-C* were physically linked, but the orientation of the genes was opposite (see Supplemental Figure 3, panel B, Additional File 1). A phylogenetic tree was constructed to examine the relationships among this 8-kb region (see Supplemental Figure 3, panel A, Additional File 1). The orthologous region from *S. bicolor* was used as an outgroup, and the separation in the two groups (Region01 and Region02) suggests that the shared 8-kb sequence appeared as the consequence of a sugarcane-specific duplication.

Region01 exhibited high gene collinearity with *S. bicolor*. However, in the BAC haplotype VII, a change in gene order involving the sorghum orthologs Sobic.003G221800 and Sobic.003G221400 was observed (Figure 1, dotted line). We were unable to determine whether this alteration resulted from a duplication or a translocation since we do not have a single haplotype that covers the entire region. Sobic.003G221800 is missing in this position from haplotypes I, II and VI.

Region01 and Region02, except for the genes *HP600* and *CENP-C*, contain different sorghum orthologous genes (Figure 1). Region02 was found to be noncollinear with *S. bicolor* (Figures 1 and 2), which reinforces the notion of a specific duplication in sugarcane. Region02 appeared as a mosaic formed by different sorghum orthologous genes distributed in different chromosomes and arose by duplication after the separation of sorghum and sugarcane.

In Region02, the Sobic.008G134300 orthologous gene was found only in haplotype VIII, and the Sobic.008G134700 ortholog was found in a different position in haplotype IX (Figure 1, dotted line in Region02 and Figure 2). The phylogenetic analysis of Sobic.008G134700 and sugarcane orthologs demonstrated that sugarcane haplotype IX are more closely related to sorghum than to other sugarcane homeologs (see Supplemental Figure 4, Additional File 1). In addition, the orientation of transcription of the Sobic.008G134700 ortholog in haplotype IX is opposite that of the other sugarcane haplotypes (Figures 1 and 2). This finding suggests that this gene could be duplicated (paralogs) or translocated (orthologs) in haplotypes X, XIV, XV and XVI. No *S. bicolor* orthologous region that originated from Region02 could be determined, since it contained genes from multiple sorghum chromosomes.

Twenty long terminal repeat (LTR) retrotransposons were located in the two regions, but no LTR retrotransposons were shared among the haplotypes from Region01 and Region02, suggesting that all LTR retrotransposon insertions occurred after the duplication. In addition, ancient LTR retrotransposons could be present, but the sequences among the sugarcane haplotypes are so divergent that they could not be identified. The oldest LTR retrotransposon insertions were dated from 2.3 Mya (from haplotype VIII from Region02, a DNA/MuDR transposon, similar to MUDR1N_SB), which means that there is evidence that this duplication is at least 2.3 Mya old. Four LTR retrotransposons similar to RLG_scAle_1_1-LTR had identical sequences (Region01: Sh083P14_TE0360 – haplotype III and Sh040F02_TE0180 – haplotype XI; Region02: Sh285K15_TE0060 – haplotype XII and Sh452C23_TE0090 – haplotype XIII), which indicates a very recent insertion into the duplication from both regions.

To estimate the genomic diversity in sugarcane haplotypes from both regions (analyzed together and separately), the shared 8-kb region (duplication) was used (see Supplemental Table 4, Additional File 1), and the SNPs were identified. The diversity in the *HP600* and *CENP-C* genes was analyzed, and one SNP was observed every 43 bases (Region02) and 70 bases (Region01). We searched for SNPs that could distinguish each region (see Supplemental Table 5, Additional File 1) in the *HP600* and *CENP-C* genes, and one SNP was found for every 56 bases (20 SNPs in total). In addition, small (3-10 bases) and large (30 – 200 bases) insertions were found. These results revealed a high level of diversity in sugarcane, i.e., a high number of SNPs in each region, which could be used to generate molecular markers and to improve genetic maps. In addition, the diversity rate of both regions together could be used as an indicator of a duplicated gene, i.e., a rate < 20 (see Supplemental Table 4, Additional File 1).

### HP600 *and* CENP-C *haplotypes and phylogenetics*

Gene haplotypes, i.e., genes with the same coding sequences (CDSs), from *HP600* and *CENP-C* that have the same coding sequence (i.e., exons) in different BAC haplotypes were considered the same gene haplotype. In Region01, four haplotypes of *HP600* were identified. In sorghum, the size of *HP600* is 187 amino acids (561 base pairs). *HP600* has two different sizes in sugarcane haplotypes of 188 amino acids (564 base pairs – haplotype I/II/VI, haplotype IV/V and haplotype VII) and 120 amino acids (360 base pairs – haplotype III). *HP600* haplotype III has a base deletion at position 77, causing a frameshift that results in a premature stop codon.

In Region02, *HP600* exhibited six haplotypes: haplotype VIII, haplotype IX, haplotype X/XI/XII/XIII/XIV, haplotype XV, and haplotype XVI. *HP600* Haplotype IX carried an insertion of eight bases in the last exon that caused a frameshift.

In *S. bicolor, CENP-C* is formed by 14 exons [33] encoding 694 amino acids (2082 base pairs). In sugarcane, the haplotypes from Region01 had 14 exons that give rise to a protein of 708 or 709 amino acids (2124 or 2127 bases). Talbert et al. [33] described two haplotypes in sugarcane EST clones [15], *CENP-C1* and *CENP-C*2, which correspond to the haplotypes I/II and haplotypes IV/V, respectively. In addition to CENP-C1 and CENP-C2, three other *CENP-C* haplotypes were observed: haplotype III, haplotype VI, and haplotype VIII.

In Region02, the sugarcane duplication of *CENP-C* consisted of the last seven exons (exons eight to fourteen from *CENP-C* in Region01), and six haplotypes were found: haplotype VIII, haplotype IX, haplotypes XI/XII/XIII, haplotype XIV, haplotype XV, and haplotype XVI. The haplotype × BAC sequence finished before the CENP-C gene (Figure 1).

To reconstruct a phylogenetic tree for *HP600* and *CENP-C* from both regions, the orthologs from *O. sativa* and *Zea mays* L. were searched. The rice *HP600* and *CENP-C* orthologs, LOC_Os01g43060 and LOC_Os01g43050, were recovered, respectively. Maize has gone through tetraploidization since its divergence from sorghum approximately 12 million years ago [37]. The maize *HP600* ortholog search returned three possible genes with high similarity: GRMZM2G114380 (chromosome 03), GRMZM2G018417 (chromosome 01) and GRMZM2G056377 (chromosome 01). The *CENP-C* maize ortholog search returned three possible genes with high similarity: GRMZM2G114315 (chromosome 03), GRMZM2G134183 (chromosome 03), and GRMZM2G369014 (chromosome 01).

Given the gene organization among the BACs, sorghum and rice revealed that *HP600* and *CENP-C* were side by side, and the expected orthologs from maize could be GRMZM2G114380 (*HP600*) and GRMZM2G114315 (*CENP-C*) because only these two genes are physically side by side. The other maize orthologs were probably maize paralogs that resulted from specific duplications of the *Z. mays* genome.

Two phylogenetic trees were constructed (see Supplemental Figure 5, Additional File 1), one for *HP600* (see Supplemental Figure 5, panel A, Additional File 1) and the other for *CENP-C* (see Supplemental Figure 5 panel B, Additional File 1), using sugarcane *HP600* and *CENP-C* haplotypes from both regions. The results demonstrated that the haplotypes from Region01 and Region02 are more similar to themselves than they are to those of sorghum or rice. Thus, the results also suggest that Region02 contains paralogous genes from Region01.

The divergence times among sugarcane *HP600* haplotypes and sorghum ranged from 1.5 Mya to 4.5 Mya. For *CENP-C*, the haplotype divergence time rates were 0.3-0.7 Mya, and the comparison with sorghum indicated 4.2-4.5 Mya for the highest values. The estimated sugarcane × sorghum divergence time was 5 Mya [38] to 8-9 Mya [29, 39].

### Chromosome number determination and BAC-FISH

The determination of the range of *CENP-C* and *HP600* loci that are present in the sugarcane genome was performed using in situ hybridization. First, the number of chromosomes in sugarcane variety SP80-3280 was defined, but the number of clear and well-spread metaphases for the variety SP80-3280 was less than 10 (see Supplemental Table 6, Additional File 1). We expanded the analysis to four more varieties of sugarcane (SP81-3250, RB835486, IACSP95-3018 and IACSP93-3046) to improve the conclusions (see Supplemental Figure 6 – Panel A-E – and Supplemental Table 6, Additional File 1). The most abundant number of chromosomes was 2n = 112 (range: 2n = 98 to 2n = 118 chromosomes). The chromosome number of the *Saccharum* hybrid cultivar SP80-3280 was found to be 2n = 112 (range: 2n = 108 to 2n = 118 chromosomes - see Supplemental Table 6, Additional File 1). Vieira et al. [40] also identified 2n = 112 chromosomes in the IACSP93-3046 variety, corroborating our data. The 2n = 112 chromosome number should indicate convergence in the number of chromosomes in the *Saccharum* hybrid cultivar.

As second step, we used two varieties with the best chromosome spreads, i.e., IACSP93-3046 and IACSP95-3018, for the CMA/DAPI banding patterns (see Supplemental Figure 6 – Panel F-I, Additional File 1). The variety IACSP93-3046 exhibited at least six terminal CMA^+^/DAPI^-^ bands, one chromosome with CMA^+^/DAPI^o^ and two chromosomes with adjacent intercalations of CMA^+^ and DAPI^+^ in the same chromosome (see Supplemental Figure 6 – Panel F and G, Additional File 1). The variety IACSP95-3018 revealed seven terminal CMA^+^/DAPI^-^ bands, and at least two chromosomes exhibited adjacent CMA^+^ and DAPI^+^; one was in the intercalary position, and the other was in the terminal position (see Supplemental Figure 6 – Panel H and I, Additional File 1). Additionally, the equal number of chromosomes and the divergent number of bands could indicate different chromosomic arrangements and/or different numbers of homeologs in each variety.

Finally, we performed BAC-FISH in the better metaphases of variety SP80-3280 using Shy064N22 (haplotype VII) from Region01; 64 metaphases with some signal of hybridization were obtained, and for the BAC-FISH of Shy048L15 (haplotype XI) from Region02, 69 were obtained. At least six metaphases for each region were used to determine the number of signals. For BAC Shy064N22 Region01, eight signals could be counted (Figure 3 – Panel A), and for BAC Shy048L15 in Region02, ten signals could be defined (Figure 3 – Panel B). These results detail the numbers of haplotypes in sugarcane for Region01 and Region 02. Moreover, the numbers of BAC haplotypes found in each region are appropriate considering the BAC-FISH results, suggesting a missing haplotype for each region.

**Figure 3.**
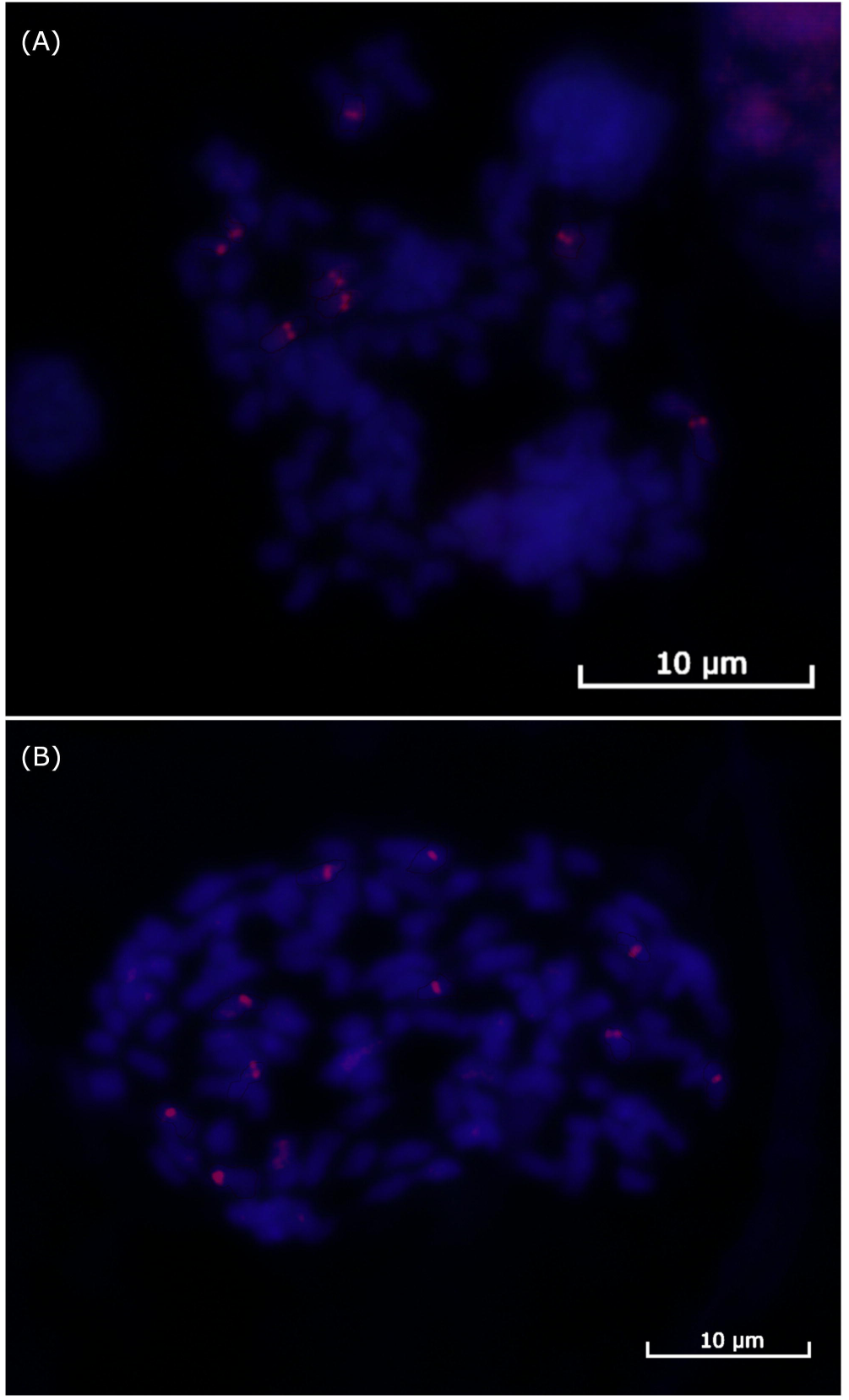
FISH hybridization of the sugarcane BACs. Panel (A): BAC Shy065N22 hybridization in sugarcane variety SP-803280 mitosis showing eight signals for Region01. Panel (B): BAC Shy048L15 hybridization in sugarcane variety SP-803280 mitosis showing ten signals for Region02.

The results observed so far suggest differences between the haplotypes, i.e., different TEs, insertions and even gene insertions/translocations. We used an identity of 99% to determinate the same BAC haplotype. The possibility of haplotypes with more than 99% similarity *in vivo* could not be tested with our data, since it is not possible distinguish a mismatch in sequence assembly from a real haplotype.

### Expression of *HP600* and *CENP-C* haplotypes

The transcriptomes of the sugarcane variety SP80-3280 from the roots, shoots and stalks were mapped on *HP600* and *CENP-C* (NCBI SRR7274987), and the set of transcripts was used for the transcription analyses. All of the haplotypes of *HP600* from Region01 were covered by the reads, including haplotype III with a premature stop codon. None of the haplotypes of *HP600* from Region02 were found, suggesting *HP600* from Region02 is not expressed (see Supplemental Figure 3, Additional File 1). For the *CENP-C* gene from Region01, the haplotypes IV/V were found to be expressed. Furthermore, haplotypes I/II, haplotype VI and haplotype VII were fully covered by the reads, except for the first three SNPs, but these SNPs were described in the work of Talbert et al. [33] under the haplotype CENP-C1, suggesting that the set of reads did not cover this region. For haplotype III, one SNP was not found, but nine exclusive SNPs of this haplotype were represented. Therefore, all haplotypes of *CENP-C* from Region01 were considered to be expressed.

The *CENP-C* haplotypes I/II, III and VI from Region01 have large retrotransposons in the introns (Figure 2 – black rectangles). Additionally, no evidence of substantial influence on expression could be found for this gene, which may indicate the silencing of these LTR retrotransposons, as discussed by Kim and Zilberman [41].

The mapping of the transcript reads in the *CENP-C* haplotypes from Region02 revealed evidence of a chimerical gene (Figure 1, dotted rectangle and Figure 4). The chimeric gene was formed by the first five exons of the sugarcane orthologous gene of Sobic.003G299500 and the eighth to fourteenth exons of *CENP-C* (Figure 4 – Panel C). RNAseq reads overlapped the region corresponding to the union of the chimerical gene (position 1253 of the *CENP-C* haplotypes from Region02 by 38 reads - Figure 4 – Panel F). This result provided robust evidence for the formation of the chimerical gene and its expression.

**Figure 4.**
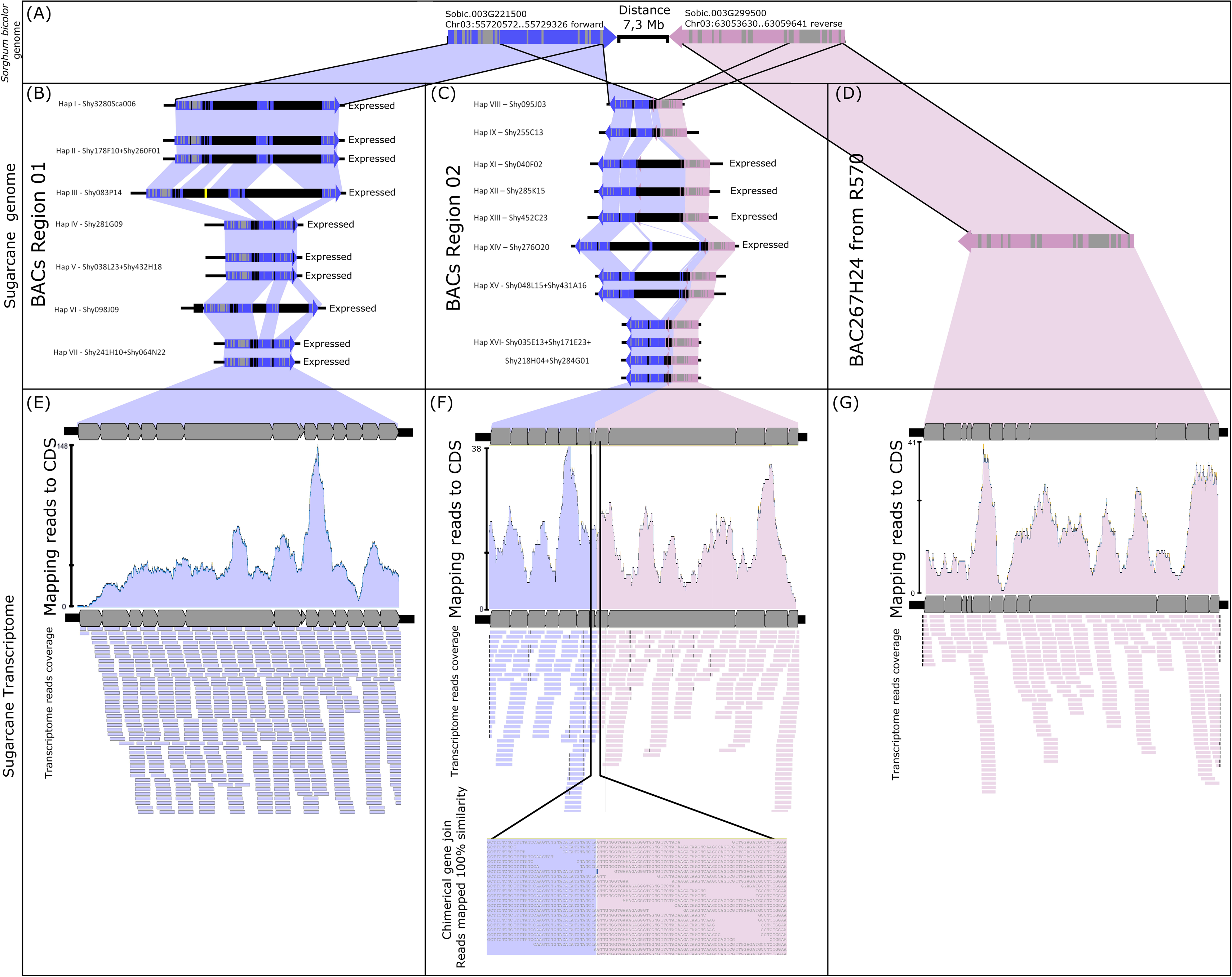
Fusion gene formation of *CENP-C* and Sobic003G299500. Panel (A): Sorghum *CENP-C* and Sobic003G299500 genome location. Panel (B): Sugarcane genomic *CENP-C* haplotypes in Region01 (all expressed). Panel (C): Partially duplicated sugarcane paralogs of *CENP-C* and Sobic003G299500 haplotypes in Region02 (only haplotypes XI/XII/XIII and haplotype XIV have evidence of expression). Panel (D): Sugarcane ortholog of Sobic003G299500 found in the sugarcane R570 BAC library. Panel (E): Transcripts from sugarcane SP80-3280 mapped against the CDS of sugarcane *CENP-C* haplotypes from Region01. Panel (F): Transcripts from sugarcane SP80-3280 mapped against the sugarcane chimerical paralogs of *CENP-C* and Sobic003G299500. As evidence of fusion gene formation, the transcripts show the fusion point of the paralogs. Panel (G): Transcripts from sugarcane SP80-3280 mapped against the CDS of the sugarcane R570 ortholog of Sobic003G299500.

The sugarcane gene orthologous to Sobic.003G299500 was represented by BAC BAC267H24 (GenBank KF184671) from the sugarcane hybrid R570 as published by De Setta et al. [28] under the name “SHCRBa_267_H24_F_10” (Figure 4 – Panel D). This finding indicated that the ancestral genes from sorghum (orthologs) were retained in the sugarcane genome (Figure 4 – Panel B and D) and that the chimerical gene was formed by the fusion of a partial duplication of *CENP-C* and the sorghum ortholog gene Sobic.003G299500 (Figure 4 – Panel C).

Two chimerical *CENP-C* haplotypes from Region02 were fully mapped with transcripts, i.e., haplotypes XI/XII/XIII and haplotype XIV. The chimerical *CENP-C* haplotypes IX and XVI from Region02 were not fully mapped, but exclusive SNPs from these haplotypes were recovered. The *CENP-C* haplotypes VIII and XV from Region02 exhibited no exclusive SNPs in the transcriptome, and evidence for the expression of these two haplotypes remains undefined.

### How locus number of homeologs influences expression

We searched the SNPs in the BAC sequences and RNAseq reads (i.e., only in the transcriptome of the sugarcane variety SP80-3280 from the roots, shoots and stalks – NCBI SRR7274987) and compared the correspondences to the genes *HP600* and *CENP-C*. For Region01 and Region02, we defined the ploidies as eight and ten, respectively, based on the BAC-FISH data. The numbers of BAC haplotypes recovered for Region01 and Region02 were seven and nine, respectively, which indicated one missing BAC haplotype in each region.

The missing BAC haplotypes were determined by searching for SNPs present only in the transcriptome. For the *HP600* haplotypes from Region01 (Table 1), six SNPs were found in the transcriptome and not in the BAC haplotypes, including a (GAG)3 -> (GAG)2 deletion. For the *CENP-C* gene (Table 2), eight SNPs were not represented in the genomic haplotypes. The presence of SNPs only in transcript data corroborates the assumption that (at least) one genomic haplotype was missing in each region.

**Table 1.**
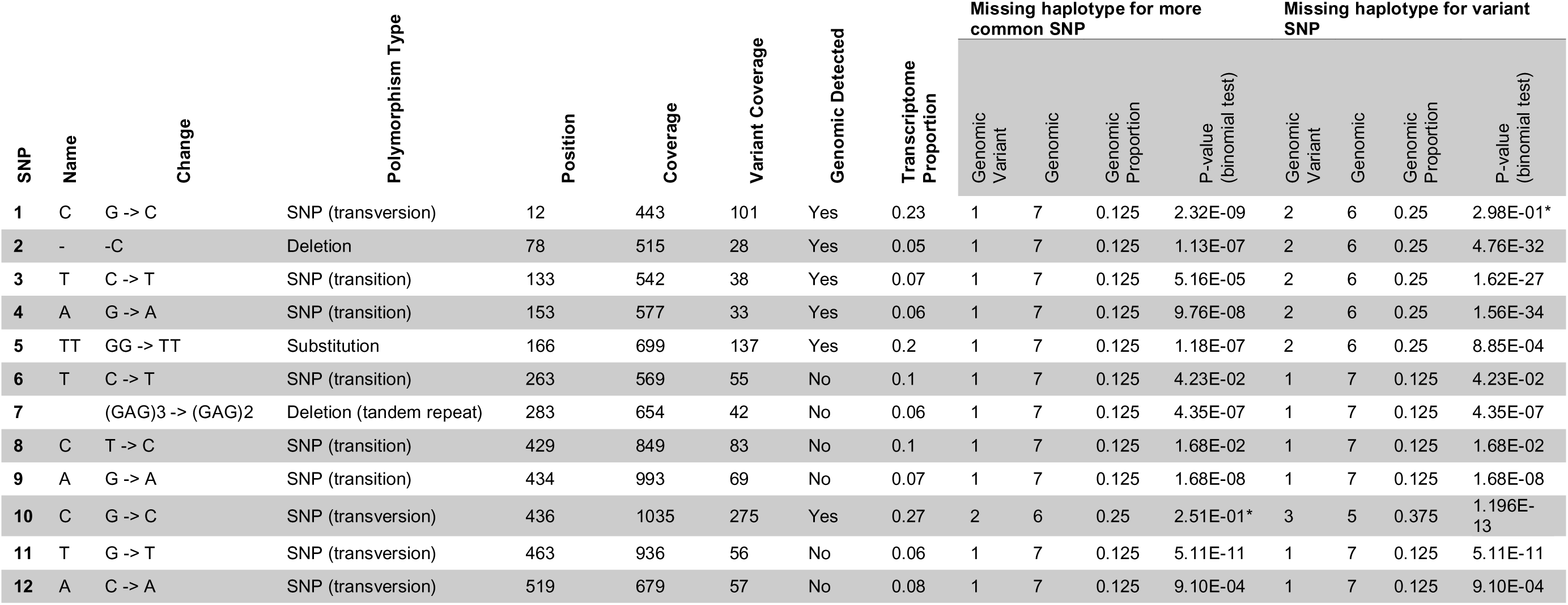
Genomic frequencies of the SNPs in the *HP600* haplotypes in Region01. Genome and transcriptome SNPs was used. The global expression (in diverse tissues) was used to determine whether the genomic frequency could explain the transcription frequency (H_0_). The binomial test was used to verify H_0_. The highlighted p-values reflect the acceptance of H_0_.

**Table 2.**
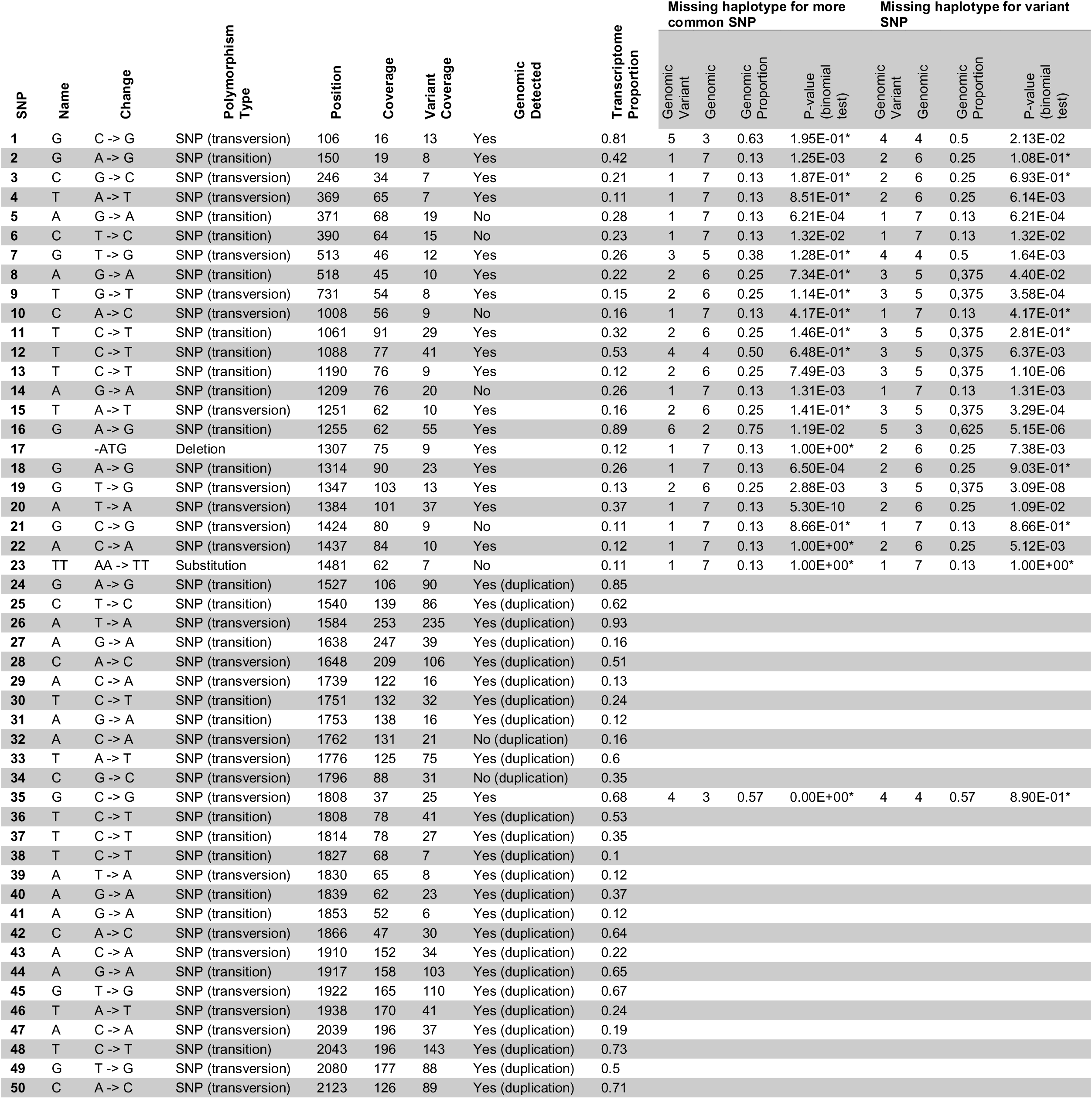
Genomic frequencies of the SNPs in the *CENP-C* haplotypes in Region01 and Region02. Genome and transcriptome SNPs was used. The global expression (in diverse tissues) was used to determine whether the genomic frequency could explain the transcription frequency (H_0_). The binomial test was used to verify H_0_. The highlighted p-values reflect the acceptance of H_0_.

| SNP | Name | Change | Polymorphism Type | Position | Coverage | Variant Coverage | Genomic Detected | Transcriptome Proportion | Missing haplotype for more common SNP |  |  |  | Missing haplotype for variant SNP |  |  |  |
| --- | --- | --- | --- | --- | --- | --- | --- | --- | --- | --- | --- | --- | --- | --- | --- | --- |
|  |  |  |  |  |  |  |  |  | Genomic Variant | Genomic | Genomic Proportion | P-value (binomial test) | Genomic Variant | Genomic | Genomic Proportion | P-value (binomial test) |
| 1 | G | C -> G | SNP (transversion) | 106 | 16 | 13 | Yes | 0.81 | 5 | 3 | 0.63 | 1.95E-01* | 4 | 4 | 0.5 | 2.13E-02 |
| 2 | G | A -> G | SNP (transition) | 150 | 19 | 8 | Yes | 0.42 | 1 | 7 | 0.13 | 1.25E-03 | 2 | 6 | 0.25 | 1.08E-01* |
| 3 | C | G -> C | SNP (transversion) | 246 | 34 | 7 | Yes | 0.21 | 1 | 7 | 0.13 | 1.87E-01* | 2 | 6 | 0.25 | 6.93E-01* |
| 4 | T | A -> T | SNP (transversion) | 369 | 65 | 7 | Yes | 0.11 | 1 | 7 | 0.13 | 8.51E-01* | 2 | 6 | 0.25 | 6.14E-03 |
| 5 | A | G -> A | SNP (transition) | 371 | 68 | 19 | No | 0.28 | 1 | 7 | 0.13 | 6.21E-04 | 1 | 7 | 0.13 | 6.21E-04 |
| 6 | C | T -> C | SNP (transition) | 390 | 64 | 15 | No | 0.23 | 1 | 7 | 0.13 | 1.32E-02 | 1 | 7 | 0.13 | 1.32E-02 |
| 7 | G | T -> G | SNP (transversion) | 513 | 46 | 12 | Yes | 0.26 | 3 | 5 | 0.38 | 1.28E-01* | 4 | 4 | 0.5 | 1.64E-03 |
| 8 | A | G -> A | SNP (transition) | 518 | 45 | 10 | Yes | 0.22 | 2 | 6 | 0.25 | 7.34E-01* | 3 | 5 | 0,375 | 4.40E-02 |
| 9 | T | G -> T | SNP (transversion) | 731 | 54 | 8 | Yes | 0.15 | 2 | 6 | 0.25 | 1.14E-01* | 3 | 5 | 0,375 | 3.58E-04 |
| 10 | C | A -> C | SNP (transversion) | 1008 | 56 | 9 | No | 0.16 | 1 | 7 | 0.13 | 4.17E-01* | 1 | 7 | 0.13 | 4.17E-01* |
| 11 | T | C -> T | SNP (transition) | 1061 | 91 | 29 | Yes | 0.32 | 2 | 6 | 0.25 | 1.46E-01* | 3 | 5 | 0,375 | 2.81E-01* |
| 12 | T | C -> T | SNP (transition) | 1088 | 77 | 41 | Yes | 0.53 | 4 | 4 | 0.50 | 6.48E-01* | 3 | 5 | 0,375 | 6.37E-03 |
| 13 | T | C -> T | SNP (transition) | 1190 | 76 | 9 | Yes | 0.12 | 2 | 6 | 0.25 | 7.49E-03 | 3 | 5 | 0,375 | 1.10E-06 |
| 14 | A | G -> A | SNP (transition) | 1209 | 76 | 20 | No | 0.26 | 1 | 7 | 0.13 | 1.31E-03 | 1 | 7 | 0.13 | 1.31E-03 |
| 15 | T | A -> T | SNP (transversion) | 1251 | 62 | 10 | Yes | 0.16 | 2 | 6 | 0.25 | 1.41E-01* | 3 | 5 | 0,375 | 3.29E-04 |
| 16 | G | A -> G | SNP (transition) | 1255 | 62 | 55 | Yes | 0.89 | 6 | 2 | 0.75 | 1.19E-02 | 5 | 3 | 0,625 | 5.15E-06 |
| 17 |  | -ATG | Deletion | 1307 | 75 | 9 | Yes | 0.12 | 1 | 7 | 0.13 | 1.00E+00* | 2 | 6 | 0.25 | 7.38E-03 |
| 18 | G | A -> G | SNP (transition) | 1314 | 90 | 23 | Yes | 0.26 | 1 | 7 | 0.13 | 6.50E-04 | 2 | 6 | 0.25 | 9.03E-01* |
| 19 | G | T -> G | SNP (transversion) | 1347 | 103 | 13 | Yes | 0.13 | 2 | 6 | 0.25 | 2.88E-03 | 3 | 5 | 0,375 | 3.09E-08 |
| 20 | A | T -> A | SNP (transversion) | 1384 | 101 | 37 | Yes | 0.37 | 1 | 7 | 0.13 | 5.30E-10 | 2 | 6 | 0.25 | 1.09E-02 |
| 21 | G | C -> G | SNP (transversion) | 1424 | 80 | 9 | No | 0.11 | 1 | 7 | 0.13 | 8.66E-01* | 1 | 7 | 0.13 | 8.66E-01* |
| 22 | A | C -> A | SNP (transversion) | 1437 | 84 | 10 | Yes | 0.12 | 1 | 7 | 0.13 | 1.00E+00* | 2 | 6 | 0.25 | 5.12E-03 |
| 23 | TT | AA -> TT | Substitution | 1481 | 62 | 7 | No | 0.11 | 1 | 7 | 0.13 | 1.00E+00* | 1 | 7 | 0.13 | 1.00E+00* |
| 24 | G | A -> G | SNP (transition) | 1527 | 106 | 90 | Yes (duplication) | 0.85 |  |  |  |  |  |  |  |  |
| 25 | C | T -> C | SNP (transition) | 1540 | 139 | 86 | Yes (duplication) | 0.62 |  |  |  |  |  |  |  |  |
| 26 | A | T -> A | SNP (transversion) | 1584 | 253 | 235 | Yes (duplication) | 0.93 |  |  |  |  |  |  |  |  |
| 27 | A | G -> A | SNP (transition) | 1638 | 247 | 39 | Yes (duplication) | 0.16 |  |  |  |  |  |  |  |  |
| 28 | C | A -> C | SNP (transversion) | 1648 | 209 | 106 | Yes (duplication) | 0.51 |  |  |  |  |  |  |  |  |
| 29 | A | C -> A | SNP (transversion) | 1739 | 122 | 16 | Yes (duplication) | 0.13 |  |  |  |  |  |  |  |  |
| 30 | T | C -> T | SNP (transition) | 1751 | 132 | 32 | Yes (duplication) | 0.24 |  |  |  |  |  |  |  |  |
| 31 | A | G -> A | SNP (transition) | 1753 | 138 | 16 | Yes (duplication) | 0.12 |  |  |  |  |  |  |  |  |
| 32 | A | C -> A | SNP (transversion) | 1762 | 131 | 21 | No (duplication) | 0.16 |  |  |  |  |  |  |  |  |
| 33 | T | A -> T | SNP (transversion) | 1776 | 125 | 75 | Yes (duplication) | 0.6 |  |  |  |  |  |  |  |  |
| 34 | C | G -> C | SNP (transversion) | 1796 | 88 | 31 | No (duplication) | 0.35 |  |  |  |  |  |  |  |  |
| 35 | G | C -> G | SNP (transversion) | 1808 | 37 | 25 | Yes | 0.68 | 4 | 3 | 0.57 | 0.00E+00* | 4 | 4 | 0.57 | 8.90E-01* |
| 36 | T | C -> T | SNP (transition) | 1808 | 78 | 41 | Yes (duplication) | 0.53 |  |  |  |  |  |  |  |  |
| 37 | T | C -> T | SNP (transition) | 1814 | 78 | 27 | Yes (duplication) | 0.35 |  |  |  |  |  |  |  |  |
| 38 | T | C -> T | SNP (transition) | 1827 | 68 | 7 | Yes (duplication) | 0.1 |  |  |  |  |  |  |  |  |
| 39 | A | T -> A | SNP (transversion) | 1830 | 65 | 8 | Yes (duplication) | 0.12 |  |  |  |  |  |  |  |  |
| 40 | A | G -> A | SNP (transition) | 1839 | 62 | 23 | Yes (duplication) | 0.37 |  |  |  |  |  |  |  |  |
| 41 | A | G -> A | SNP (transition) | 1853 | 52 | 6 | Yes (duplication) | 0.12 |  |  |  |  |  |  |  |  |
| 42 | C | A -> C | SNP (transversion) | 1866 | 47 | 30 | Yes (duplication) | 0.64 |  |  |  |  |  |  |  |  |
| 43 | A | C -> A | SNP (transversion) | 1910 | 152 | 34 | Yes (duplication) | 0.22 |  |  |  |  |  |  |  |  |
| 44 | A | G -> A | SNP (transition) | 1917 | 158 | 103 | Yes (duplication) | 0.65 |  |  |  |  |  |  |  |  |
| 45 | G | T -> G | SNP (transversion) | 1922 | 165 | 110 | Yes (duplication) | 0.67 |  |  |  |  |  |  |  |  |
| 46 | T | A -> T | SNP (transversion) | 1938 | 170 | 41 | Yes (duplication) | 0.24 |  |  |  |  |  |  |  |  |
| 47 | A | C -> A | SNP (transversion) | 2039 | 196 | 37 | Yes (duplication) | 0.19 |  |  |  |  |  |  |  |  |
| 48 | T | C -> T | SNP (transition) | 2043 | 196 | 143 | Yes (duplication) | 0.73 |  |  |  |  |  |  |  |  |
| 49 | G | T -> G | SNP (transversion) | 2080 | 177 | 88 | Yes (duplication) | 0.5 |  |  |  |  |  |  |  |  |
| 50 | C | A -> C | SNP (transversion) | 2123 | 126 | 89 | Yes (duplication) | 0.71 |  |  |  |  |  |  |  |  |

Using the results obtained from the RNAseq mapping of haplotypes, we also assumed that all haplotypes of the gene *HP600* were expressed in Region01 and that none were expressed in Region02. For *CENP-C*, all haplotypes from Region01 were considered expressed, and it was not possible to identify how many haplotypes were expressed in Region02 (chimerical gene); thus, we used only the nonduplicated portion of *CENP-C* (exons one to seven of the *CENP-C* gene).

We formed three assumptions using the previous results: (I) there is a missing haplotype for each region; (II) all haplotypes of HP600 from Region01 are expressed, and there is no expression of *HP600* in Region02; and (III) *CENP-C* is expressed in both regions, but only in Region01 is it possible to infer that all haplotypes are expressed. Using these premises, we investigated the possibilities of one BAC haplotype being expressed at a higher or lower level than the others. Therefore, if the haplotypes contribute equally to expression, one SNP found in a BAC should have the same ratio (dosage) for the transcriptome data. Since we found evidence for a missing haplotype, two tests were performed: (I) we determined whether the missing BAC haplotype contributed to the dosage of more common SNPs, and (II) we determined whether the missing BAC haplotype contributed to the dosage of the variant SNP.

For the *HP600* haplotypes from Region01 (Table 1), only the SNPs 10 and 1 had significant p-values for hypotheses (I) and (II), respectively. These results suggested that the BAC haplotype ratio does not explain the transcriptome ratio. The transcript frequencies of SNPs 2, 3, and 4 (Table 1) were less than 0.125 (the minimum expected ratio for 1:7). To explain these frequencies, the dosage of the SNPs should be higher than a ploidy of eight (i.e., more than twelve), and our data do not support this possibility. The three variant SNPs came from *HP600* haplotype III. This finding could be evidence of some differential expression of the gene haplotypes, which could suggest that haplotype III is expressed at a lower level than the others for the HP600 gene.

For *CENP-C*, only the nonduplicated portions of the haplotypes from Region01 were used. At least one hypothesis was accepted for 17 (70%) SNPs (Table 2). The mean coverage of the SNPs was 64 reads per SNP, which could be considered a low coverage when an eight-ploidy region (Region01) is being inspected (Table 2). Moreover, the result suggests that the haplotypes from Region01 are equally expressed.

### Genetic mapping

For the genetic mapping, 44 SNPs (see Supplemental Table 7, Additional File 1) were used to develop molecular markers (Figure 5), and they were used to construct a genetic map. The SuperMASSA [42] software calculates all possible ploidies for a locus and produces the most probable ploidy. Moreover, it is possible to define a fixed ploidy for a locus. The first option was used to call the dosages, which were ultimately used to construct the genetic map (Figure 6), and this map was compared with the fixed ploidy according to the BAC haplotype results (Figure 5). In fact, when using a Bayesian approach similar to that from the SuperMASSA, providing prior information about the ploidy level might improve the dosage estimates.

**Figure 5.**
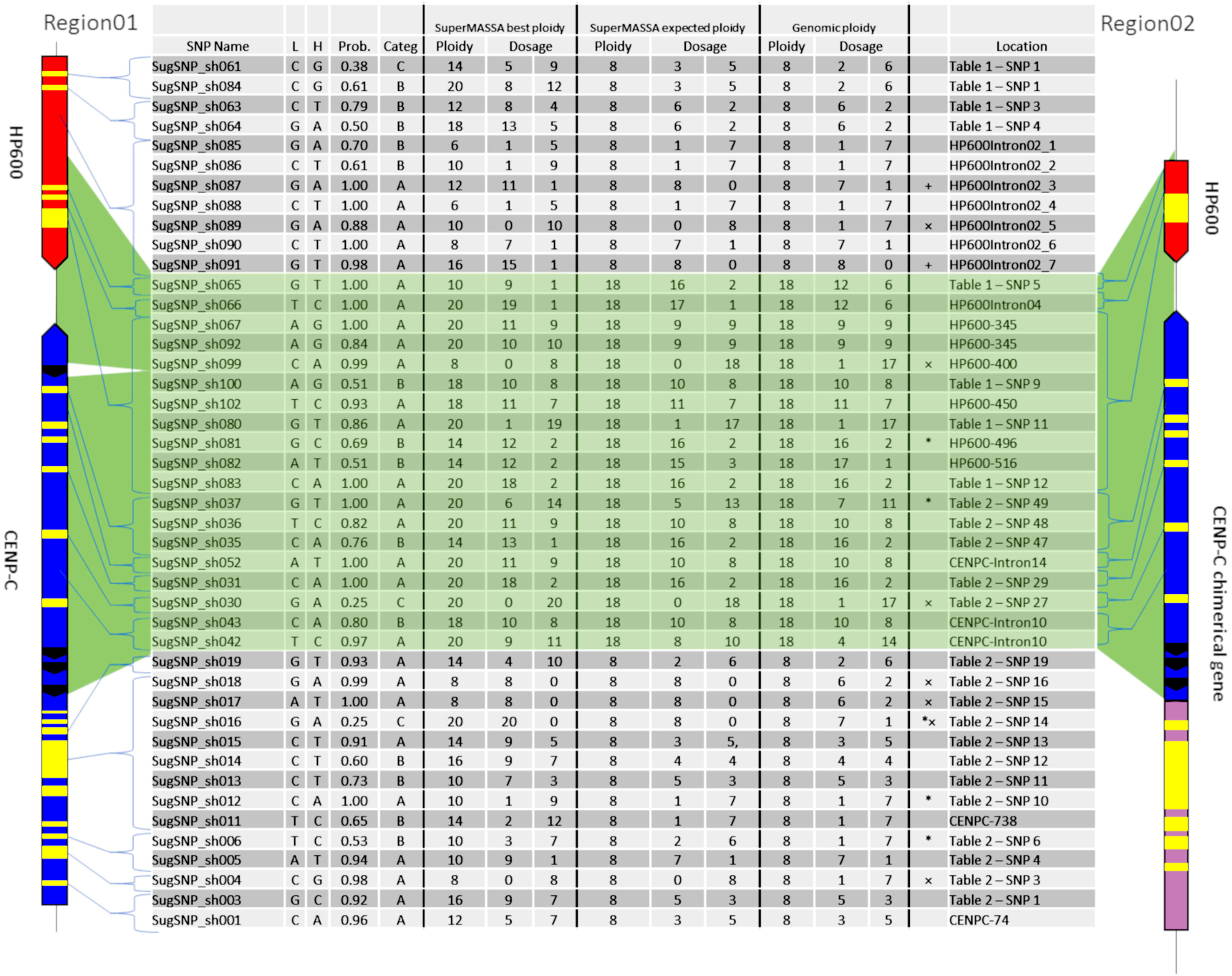
Ploidy and dosage in the sugarcane genomic DNA (BACs) and SuperMASSA estimation. The location of each SNP is shown by one haplotype from Region01 and one haplotype from Region02. “SuperMASSA Best Ploidy” means the SuperMASSA best ploidy with a posteriori probability of >0.8. “SuperMASSA Expected Ploidy” means we fixed the ploidy of the loci in SuperMASSA according to the BAC-FISH and BAC sequencing results. “Genomic Ploidy” means the ploidy of the loci according to the BAC-FISH and BAC sequencing results. “*” means the SNP was found only in the transcriptome.

**Figure 6.**
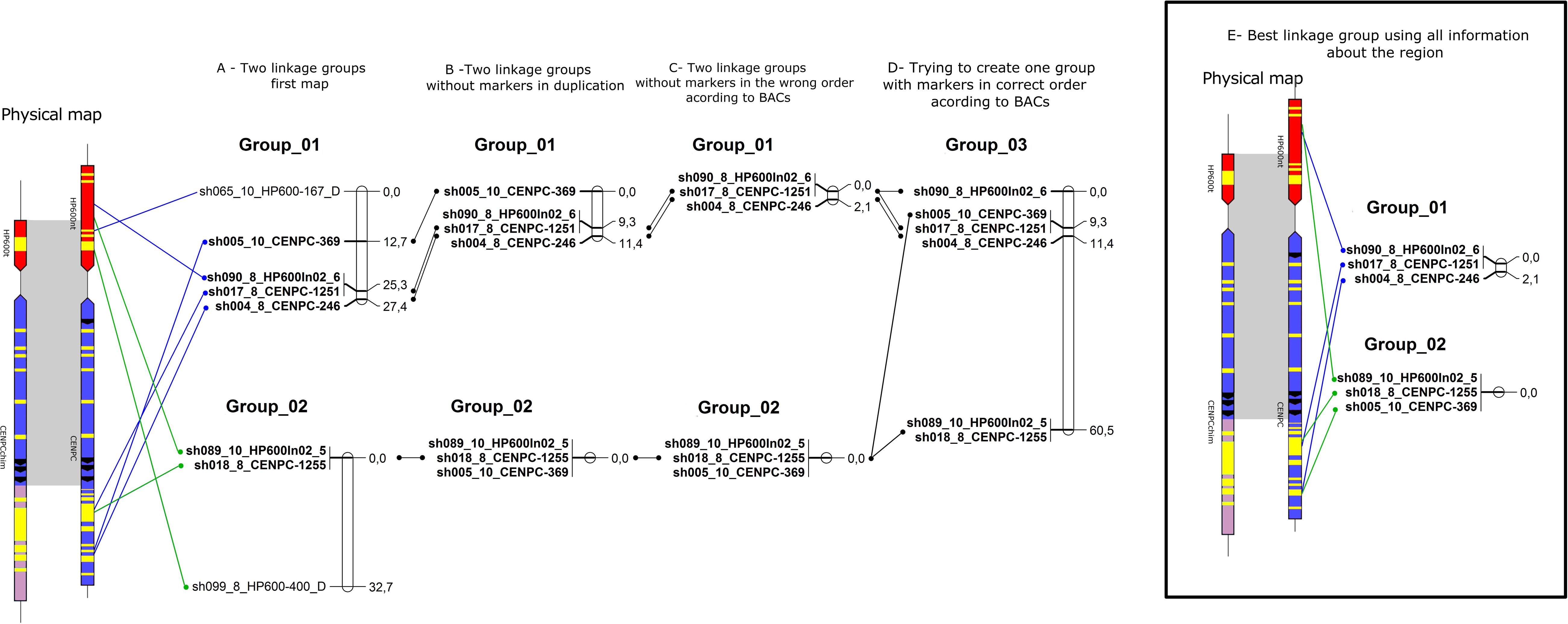
Schematic representation of sugarcane linkage map. The sugarcane variety SP80-3280 SNPs was used to create multiples linkage map with information about the sugarcane genome (BACs).

The markers from introns and exons were drawn along Region01 (Figure 5, “Location” column), including the duplicated region found in Region02. Among them, seven exhibited no variant presence in genotyping (Figure 5 – “×” marked), but five were detected in the RNAseq reads. Two markers (Figure 5 – “+” marked) were detected only for the “SuperMASSA best ploidy”, which was a ploidy higher than the “SuperMASSA expected ploidy”. In addition, two SNP loci were genotyped two times using different capture primer pairs (SugSNP_sh061/SugSNP_sh084 and SugSNP_sh067/SugSNP_sh092), and, as expected, at higher ploidy levels (> 12), the dosages of the loci diverge. These results could be explained by intrinsic problems in molecular biology that occur during the preparation of the samples, which affects the signal intensity of the Sequenom iPLEX MassARRAY® (Sequenom Inc., San Diego, CA, USA) data.

The SuperMASSA best ploidy was equal to the genomic ploidy for six SNPs (Figure 5), and the allelic dosage confirmed in four of them. When the ploidy for the loci was fixed (8 for Region01 and 18 for Region01 and Region02 SNPs), 24 SNPs had their dosage confirmed by SuperMASSA (Figure 5 – “SuperMASSA expected ploidy” columns). Notably, the estimation of the ploidy could also be a difficult task [43], but when the ploidy used was found in BAC-FISH, the estimated dosage was in agreement with the dosage found in the BACs in 63% (28) of the SNPs (Figure 5).

For the genetic mapping, ten markers were used according to the SuperMASSA best ploidy results. First, attempts were made to add each marker to the existing linkage groups published by Balsalobre et al. [21], but none of the markers could be linked to the groups. Then, the markers were tested for linkage among themselves. Two linkage groups could be created (Figure 6 – panel A) with 27.4 cM and 32.7 cM, respectively. The two linkage groups were too large; therefore, the markers SugSNP_sh065 and SugSNP_sh099 were excluded, since both markers were in the duplicated region (Figure 6 – panel B).

Then, attempts were made to add the remaining markers to the groups again, and the marker SugSNP_sh005 was inserted into Linkage group 02 (Figure 6 – panel C). The markers that were in the wrong positions according to the physical map (BACs) were also excluded, and the marker SugSNP_sh005 was excluded from Linkage group 01 but remained in Linkage group 02 (Figure 6 – panel C). Then, an attempt was made to form one linkage group with the remaining markers by forcing OneMap to place the markers in a single group. Again, the size of the group was too large (60.3 cM - Figure 6 – panel D). Therefore, the best representation of the region was two linkage groups, Linkage group 01 with 2.1 cM, and Linkage group 02 with 0 cM (Figure 6 – panel E).

## Discussion

The genetic, genomic and transcriptome interactions among sugarcane homeologs remain obscure. Several works have attempted to understand these interactions [24, 27-31, 43-47]; and others. The high polyploidy in sugarcane cultivars make the detection of the ploidy of a locus a challenge [43, 45-47]; and other.

The chromosome number of the main Brazilian varieties was determined. The chromosome number determination showed an equal number of chromosomes (2n = 112, range: 2n = 98-118). The aneuploid nature of sugarcane hybrid cultivars [9, 48] means that they contain different numbers of homeologous chromosomes. A number of differences in the CMA/DAPI patterns were found among the different varieties analyzed in this study, suggesting differences in chromosome content, i.e., differences in homeologous arrangement. Vieira et al. [40] analyzed several sugarcane pollen cells showing metaphase chromosomes that did not line up at the plate, lagging chromosomes and chromosomal bridges, tetrad cells with micronuclei and dyads with asynchronous behavior. They concluded that the presence of chromatin bridges indirectly indicates the occurrence of chromosomal inversions.

For genetic and genomic studies, information about genomic organization is very important. Here, we report the construction of two new BAC libraries for two important Brazilian cultivars, SP80-3280 and SP93-3046, with a larger number of clones and higher sugarcane genome coverage than previously reported [25-27]. The number of clones in a library is directly related to the number of homeologous regions that can be recovered.

The approach of mapping the BES in the *S. bicolor* genome, already performed for other libraries [27, 49, 50], revealed high synteny with the *S. bicolor* genome and a large number of TEs in the sugarcane genome. Kim et al. [49] showed BES anchorage of approximately 6.4%, and Figueira et al. [27] showed anchorage of approximately 22%. Our data showed 10% BES anchorage in the sorghum genome for both libraries constructed. These results are more similar to those of Kim et al. [49], since they used only BES >= 300 bp, and we used BES >= 100 bp.

The sugarcane genome has been reported to be composed of approximately 40% TEs [27, 28, 49]. We also found that the average percentage of TEs was 40%, but this value has a very large variance among the haplotypes, with a minimum of 21% and a maximum of 65%. Figueira et al. [27] and De Setta et al. [28] also revealed an inflation of the sugarcane genome in comparison with the *S. bicolor* genome. De Setta et al. [28] reported a very significant expansion that mainly occurred in the intergenic and intronic regions and was primarily because of the presence of TE, and we confirmed this report by comparing our data with data on the *S. bicolor* genome. Several studies have reported a very significant sugarcane genome expansion [24, 27-31, 44].

A hypothetical gene *HP600* and the *CENP-C* gene were used in this work as a case study. The function of *HP600* is unknown, but the ortholog of this gene is present in the genomes of rice (LOC_Os01g43060), maize (GRMZM2G114380) and sorghum (Sobic.003G221600). Sobic.003G221600 (ortholog of *HP600*) was also found in a QTL for BRIX (sugar accumulation) that was mapped by Murray et al. [51] and based on the sorghum consensus map reported by Mace and Jordan [52]. The *CENP-C* protein is a kinetochore component [35, 36] located next to *HP600*. Here, we have demonstrated the existence of paralogous genes for *HP600* and *CENP-C* that are localized in two different homeologous sugarcane chromosome groups. The BAC haplotypes could be separated into two sugarcane homeologous groups as follows: Region01 contained the collinearity region between sorghum and sugarcane *HP600* and *CENP-C* genes, and Region02 contained their paralogs.

Region01 is a recurrent case of high gene conservation and collinearity among sugarcane homeologs and the *S. bicolor* genome as reported by other authors [24, 28-31]. Region02 contains parts of the genes *HP600* and *CENP-C* (paralogs). No synteny was found between the sugarcane Region02 and the sorghum genome. In Region02, a third partial gene (ortholog of Sobic.003G299500) was also found next to *CENP-C*, and transcriptome analysis revealed the fusion of the *CENP-C* partial exons with the partial exons of the sugarcane ortholog of Sobic.003G299500 to form a chimerical gene. Region02 is a scrambled sugarcane sequence that was possibly formed from different noncollinear ancestral sequences, but the exonic structure of the genes was retained. The phylogenetic analysis of gene haplotypes from *HP600* and *CENP-C* provided evidence that the multiple genes found in maize are the result of specific duplications in the maize taxa, as expected.

The nature of sugarcane hybrid cultivars, especially the processes of polyploidization [1, 2] and nobilization [3], are the main reason for the genomic variability, gene pseudogenization and increases in new genes [10]. It is possible that the structure found in Region02 could be a result of the polyploidization and domestication of sugarcane [6, 9, 48, 53]. However, the presence of a set of sugarcane homeologs with very similar gene structures leads us to speculate that the occurrence of an ancestral event prior to polyploidization resulted in this structure.

Rearrangement events can also be caused by TEs, but they are normally caused by the formation of a pseudogene [54, 55]. In the case of Region02, multiple events resulted in this region, but the number and types (TE, translocations) of events could not be determined with our data.

BAC-FISH hybridization was used to indicate the ploidy of each region. Eight signals were found for Region01 and 10 for Region02. These results are highly consistent with the BAC haplotype and suggest that at least one BAC haplotype is missing in each region. Casu et al. [45], Xue et al. [46] and Sun and Joyce [47] reported different methods to quantify the copy number of endogenous gene, some of which resulted in odd copy numbers. Sun and Joyce [47] reported that the low or odd numbers could be explained by the contribution of only the *S. spontaneum* or the *S. officinarum* genome. The presence of genes without collinearity among the sugarcane homeologs could also explain the result as verified for the orthologs Sobic.003G221800 and Sobic.008G134700.

The genomic SNP variation in sugarcane coding regions has been estimated to be one SNP every 50 bp [56] and one every 86 bp [16]. For the coding Region01 one SNP was found per 70 bases. Feltus et al. [57] showed that different ratios of SNPs occur across the genome. When we compared Region01 and Region02 one SNP was found per 12 bases using only the data for one sugarcane variety SP-803280. In light of the possible existence of at least one more haplotype, this number could be underestimated.

Once established, the polyploidy might now fuel evolution by virtue of its polyploid-specific advantages. Vegetative propagation can lead to the retention of genes. Meiosis may or may not play a role in either the origin or maintenance of a polyploid lineage [58]. Vegetative propagation is widely used to propagate sugarcane (even for nondomesticated sugarcanes) and could explain the high variation in sugarcane and the high level of gene retention. However, it is not the only factor, with sugarcane polyploidization and nobilization also contributing to these effects.

The homologous gene expression in polyploids can be affected in different ways, i.e., the homologous genes may retain their original function, one or more copies may be silenced, or the genes may diversify in function or expression [59-62]. In complex polyploids, the roles of ploidy and genome composition in possible changes in gene expression are poorly understood [63]. Even in diploid organisms, this task is difficult, since different interactions can affect the expression of a gene, and not all homologs are guaranteed to contribute to a function [12]. The expression of the *HP600* and *CENP-C* haplotypes in Region01 could be confirmed. In Region02, the haplotypes of *HP600* were not found in the transcriptome dataset [16, 18], but at least two haplotypes of the gene *CENP-C* were expressed.

The gene haplotypes of *HP600* from Region01 exhibited unbalanced expression; i.e., for some reason, some haplotypes were expressed at greater levels than others. These findings could mean that apart from the duplication, *HP600* might be expressed as a single-copy gene wherein only the haplotypes of the *HP600* in Region01 were expressed. In addition, we could not identify the mechanisms contributing to the unbalanced expression. Therefore, the transcripts from different tissues make us speculate that some kind of tissue-specific expression could be occurring.

Numerous molecular mechanisms are involved in the creation of new genes, such as exon shuffling, retrotransposons and gene duplications (reviewed in Long et al. [64]). Gene fusions allow the physical coupling of functions, and their occurrence in the genome increases with the genome size [65]. Sandmann et al. [36] describes the function of the protein KNL2, which uses *CENPC-k* motifs to bind DNA sequence independently and interacts with the centromeric transcripts. The *CENPC* motif is characteristic of *CENP-C*. The *CENPC* motif of the rat *CENP-C* protein can bind directly to a chimeric H3/cenH3 nucleosome *in vitro* suggesting that this motif binds to cenH3 nucleosomes *in vivo*. Consequently, it is involved directly in cell division [35, 36]. The CENPC motifs described by Sandmann et al. [36], were compared with those of *CENP-C* genes in *A. thaliana, O. sativa, Z. mays* and *S. bicolor* (see Supplemental Figure 7, Additional File 1). The *CENP-C* haplotypes from Region02 (chimerical gene) have the same *CENPC* motif as that in sorghum. The *CENP-C* haplotypes from Region01 have one variation in the second residue of the *CENPC* motif, which is a glycine in sorghum and a valine in *CENP-C* haplotypes from Region01. This result suggests that the chimerical gene retained the ancestral residue at this site, whereas a mutation occurred in *CENP-C* haplotypes from Region01. Therefore, the mutation could have occurred in sorghum and in the haplotypes from Region02, but this is unlikely. This result suggests that the *CENP-C* haplotypes from Region01 and Region02 are able to bind to cenH3 nucleosomes.

The presence of the motif in the *CENP-C* haplotypes from the Region02 proteins could indicate a chimerical protein with a similar function, specific to sugarcane, that is involved in the organization of centromeric regions. Moreover, the presence of large LTR retrotransposons in the intronic region of the *CENP-C* haplotypes in Region01 does not influence the gene expression. Furthermore, two studies [66, 67] identified the inactivation of the same gene, IBM2/ANTI-SILENCING 1 (ASI1), which causes gene transcripts with methylated intronic transposons that terminate within the elements. The complete mechanisms that control LTR retrotransposon methylation require further clarification [41].

These results have several implications for the integration of transcriptome data and genomic data. First, for example, a gene such as *HP600* that demonstrates single-copy behavior in the transcriptome data and the genomic behavior of a duplicated gene can cause bias in genetic mapping. Second, a chimerical gene such as the *CENP-C* haplotypes in Region02 can result in different levels of expression of the duplicated and nonduplicated gene regions in the transcriptome data. Using the *CENP-C* gene as an example, if the gene expression quantification probe recovers the nonduplicated portion of the *CENP-C* gene, it will give an expression level only for the *CENP-C* haplotypes in Region01. In contrast, this probe quantifies the duplicated region of *CENP-C*, it will result in the quantification of *CENP-C* from both Region01 and Region02 and thus overestimate the expression of *CENP-C*. Consequently, analyses of the expression of the gene for functional studies for evaluating the balance of gene expression will be biased.

The SNPs were also used to compare the ploidy found in BACs with the results of SuperMASSA software [43]. SuperMASSA uses segregation ratios to estimate ploidy, which is not the same as estimating ploidy by chromosome counting because of the differences in estimation and the real ploidy visualized. The SNPs present in a duplication were mapped in a linkage group and demonstrated a high distance between the markers in the linkage map. The size of a genetic map is a function of the recombination fraction, so two factors influence the map size: (I) the number of recombinations found between two markers, and (II) genotyping errors. In this case, the mapping of duplicated markers is an error and is interpreted by OneMap in a recombination fraction, which inflates the map.

Two markers classified with a ploidy of 10 and one with a ploidy of 8 formed the linkage group 02. The ploidy is not a determinant for the OneMap construction of a linkage group, but the recombination fraction is. In other words, recombination fractions can still be computed between single-dose markers classified in different ploidy levels. In fact, most of nulliplex, simplex and duplex individuals will have the same dosage call using either 8 or 10 as the ploidy level. In addition, the genome data (BACs and BAC-FISH) demonstrated that all markers had the same ploidy of eight and that the physical distances among the markers were too small and thus probably resulted in the lack of recombination. The fact that we obtained two linkage groups can be explained by the possibility that single-dose markers may be linked in repulsion, and insufficient information is available to assemble all of the markers in one group. Trying to calculate the recombination fraction between markers D1 and D2 (according to the nomenclature of Wu et al. [68]) in diploids presents the same obstacle.

We observed the relationship between a linkage map and the physical map of a region in sugarcane. Indeed, it is a small region to observe whereas sugarcane has a large genome, and a linkage map is constructed based on the recombination fraction. However, it was possible to observe what happens in the genetic map when a duplicated locus was mapped.

The combination of divergent genomes within a hybrid can lead to immediate, profound and highly varied genome modifications, which could include chromosomal and molecular structural modifications [69-72] as well as epigenetic changes [73] and global transcriptomic changes [61, 62]. The integration of the genetic, genomic and transcriptomic data was used to explain the interaction of the two regions in sugarcane. *HP600* is a hypothetical gene that is next to the *CENP-C* gene, a kinetochore component responsible for the initiation of nucleosomes. The sugarcane gene haplotypes of *HP600* in Region01 and the *CENP-C* haplotypes in Region01 were duplicated in another group of homeologous chromosomes. The duplication of the *HP600* haplotypes in Region01 resulted in a paralog pseudogene in the *HP600* haplotypes in Region02. The duplication of *CENP-C* in the haplotypes of Region02 resulted in fusion with another gene, which contained the first five exons of the orthologous gene Sobic.003G299500 and exons eight to fourteen of *CENP-C*. The region where this duplication was inserted (Region02) contained at least three more genes that probably arose due to duplication, which indicates that multiple duplication events occurred in this region.

The *HP600* and *CENP-C* duplication described in this work occurred sometime after the separation of sugarcane and sorghum and before the polyploidization of the *Saccharum* genus. This result is supported by the following information: (I) the molecular clock time, (II) the genes are present in a homeologous group of chromosomes; and (III) the *CENP-k* motifs of the *CENP-C* haplotypes in Region02 are more similar to sorghum than to its paralog in sugarcane. The formation of a chimeric gene and the scrambled Region02 exhibited a specific moment of formation before *Saccharum* polyploidization, which makes us wonder which genomic event could be the result this formation. TEs carrying this region could not be found. It is also possible that TEs were inserted in this region, and the TE sequences were subsequently lost. An event that resulted in some genome instability could also be a reason. Additionally, multiple events could also have occurred.

The transcripts from SP80-3280 revealed full expression of the haplotypes of *HP600* in Region01 (in an unbalanced manner) and the lack of expression of the haplotypes *HP600* in Region02. The expression of the *HP600* haplotypes in Region01 can be considered a single-copy gene, despite the presence of the duplication. The *CENP-C* gene can be considered fully expressed, despite the low coverage of the transcriptome data. The *CENP-C* haplotypes in Region02 have four haplotypes that are considered expressed.

Currently, only markers with low dosages can be used to construct the genetic map in sugarcane, which is a limitation of the mapping method in polyploids. We attempted to map a duplicated region, which is a difficult task even for diploid organisms. Again, it is important to observe that we used a sugarcane variety with asexual reproduction and performed the genetic mapping in artificial progeny. We have no idea how the progeny genome responded to the cross, since sugarcane is aneuploid. In addition, the premise that each individual of the progeny did not miss any chromosome in the cross (aneuploidy) and the ploidy of a locus is the same in both the parents and all individuals in the progeny could be biologically untruthful. The genetic mapping demonstrates that there are obstacles that still need to be overcome in the genetic mapping of complex polyploids.

## Conclusion

This study sheds light on the influence of the genome arrangement for transcriptome and genetic map analyses in the sugarcane polyploid genome. The integration of genomic sequence arrangements, transcription profiles, cytogenetic organization and the genetic mapping approach might help to elucidate the behavior of gene expression, the genetic structure and successful sequence assembly of the sugarcane genome. Such integrated studies will undoubtedly help to enhance our understanding of complex polyploid genomes including the sugarcane genome.

Particular emphasis should be given to the determination studies of the ploidy level and of the duplication loci with the intention of better understanding complex polyploids. Such studies remain the most original and challenging in terms of understanding the sugarcane genome. From this perspective, this work presents an integrated approach to elucidate the allelic dynamics in polyploid genomes.

## Methods

### Plant material

The sugarcane varieties SP80-3280 and SPIAC93-3046 were collected from germplasm at the active site located in the Agronomic Institute of Campinas (IAC) Sugarcane Center in Ribeirão Preto, São Paulo, Brazil. The leaves were collected on dry ice and stored at −80ºC until use.

### BAC library construction and BAC-end analyses

The high-molecular-weight (HMW) DNA was prepared from the leaves as described by Peterson et al. [74] with modifications as described by Gonthier et al. [75]. The HMW DNA was embedded in low melt agarose (Lonza InCert™ Agarose, Lonza Rockland Inc., Rockland, ME, USA) and partially digested with HindIII (New England Biolabs, Ipswich, MA, USA). Next, two size selection steps were performed by pulsed field gel electrophoresis (PFGE) with a Bio-Rad CHEF Mapper system (Bio-Rad Laboratories, Hercules, CA, USA), and the selected DNA was ligated into the pIndigoBAC-5 HindIII-Cloning Ready vector (Epicenter Biotechnologies, Madison, WI, USA) as described by Chalhoub et al. [76]. The insert size was verified by preparing DNA BACs with the NucleoSpin® 96 Plasmid Core Kit (MACHEREY-NAGEL GmbH & Co., Düren, Germany), according to the kit instructions, and the DNA was digested by the NotI (New England Biolabs, Ipswich, MA, USA) restriction enzyme and analyzed by PFGE.

For the BES, 384 random BAC DNAs from each library were prepared with the NucleoSpin® 96 Plasmid Core Kit (MACHEREY-NAGEL GmbH & Co., Düren, Germany), according to the kit instructions. The sequencing reactions were performed according to the manufacturer’s instructions for the BigDye Terminator Kit (Applied Biosystems, Foster City, CA, USA). The primers used in the reactions were T7 Forward (5’ TAATACGACTCACTATAGG 3’) and M13 Reverse (5’ AACAGCTATGACCATG 3’). The PCR conditions were 95°C for 1 min followed by 90 cycles of 20 sec at 95°C, 20 sec at 50ºC and 4 min at 60°C. The samples were loaded on a 3730xl DNA Analyzer (Applied Biosystems). Sequence trimming was conducted by processing the traces using the base-calling software PHRED [77, 78], and reads with phred score < 20 were trimmed. The sequences were compared by using BLASTN in the *S. bicolor* genome from Phytozome v10.1 [79]. Only clones with forward and reverse sequence maps in the *S. bicolor* genome, with a maximum distance of 600 kb and with no hits with repetitive elements, were used to anchor the *S. bicolor* genome.

### Target gene determination

Transcripts of *S. bicolor, Z. mays* and *O. sativa* were obtained from Phytozome v10.1 [79]. Each transcript was queried against itself, and orthologous genes that resulted in redundant sequences were eliminated. From the remaining genes, the gene Sobic.003G221600 (*Sorghum bicolor* v3.1.1 – Phytozome v. 12) was chosen because it was inserted in a QTL for Brix from a study by Murray et al. [51], which identified the QTL in the SB-03 genome (*S. bicolor* v3.1.1 – Phytozome v. 12). The sequence of the gene Sobic.003G221600 was then used as query in the SUCEST-FUN database (http://sucest-fun.org/ - [15]) and the transcriptome obtained by Cardoso-Silva et al. [16] to recover sugarcane transcripts. All the transcripts obtained were aligned (MAFFT; [80]) to generate phylogenetic trees by the maximum likelihood method (PhylML 3.0; [81]).

The sugarcane transcripts were split into exons according to their annotation in *S. bicolor, Z. mays* and *O. sativa,* and exon five was used to design the probe to screen both BAC libraries (F: 5’ ATCTGCTTCTTGGTGTTGCTG 3’, R: 5’ GTCAGACACGATAGGTTTGTC 3’). DNA fragments were PCR-amplified from sugarcane SP80-3280 and SPIAC93-3046 genomic DNA with specific primers targeting the gene Sobic.003G221600. The PCR amplification conditions were 95°C for 8 min; 30 cycles of 20 sec denaturation at 95°C, 20 sec of annealing at 60ºC, and a 40 sec extension at 72°C; and a final 10 min extension at 72°C. The probes were sequenced before the screening of the BAC library.

### BAC library screening

Both BAC libraries were spotted onto high-density colony filters with the workstation QPix2 XT (Molecular Devices, Sunnyvale, CA, USA). The BAC clones were spotted in duplicate using a 7×7 pattern onto 22 × 22 cm Immobilon-Ny+ filters (Molecular Devices). The whole BAC library from the SP80-3280 sugarcane variety was spotted in four sets of filters, each one with 55 296 clones in duplicate and the whole BAC library from SPIAC93-3046 sugarcane variety was spotted in three sets of filters each with 55,296 clones in duplicate. The filters were processed as described by Roselli et al. [82]. Probe radiolabeling and filter hybridization were performed as described in Gonthier et al. [75].

The SP80-3280 BAC library was used to construct a 3D pool. A total of 110,592 clones were pooled into 12 superpools following the protocol used by Paux et al. [83]. The positive BAC clones from the SP80-3280 library were isolated, and one isolated clone was validated by qPCR. The insert size of each BAC was estimated by using an electrophoretic profile of NotI-digested BAC DNA fragments and observed by PFGE (CHEF-DRIII system, Bio-Rad) in a 1% agarose gel in 0.5× TBE buffer under the conditions described in Paiva et al. [84].

### Sequencing and assembly

Twenty-two positive BAC clones were sequenced in pools of 10 clones. One microgram of each BAC clone was used to prepare individual tagged libraries with the GS FLX Titanium Rapid Library Preparation Kit (Roche, Branford, CT, USA). BAC inserts were sequenced by pyrosequencing with a Roche GS FLX Life Sciences instrument (Branford, CT, USA) in CNRGV, Toulouse, France.

The sequences were trimmed with PHRED, vector pIndigoBAC-5 sequences and the *Escherichia coli* str. K12 substr. DH10B complete genome was masked using CROSS_MATCH, and the sequences were assembled with PHRAP [85-87] as described by De Setta et al. [28]. A BLASTN with the draft genome [23] was performed. A search was performed in the NCBI databank to find sugarcane BACs that could possibly have the target gene *HP600*.

### Sequence analysis and gene annotation

All the BACs were aligned to verify the presence of redundant sequences of homeologs. BAC clones with more than 99% similarity were considered the same homeolog. BACs that represented the same homeologs were not combined. The BACs were annotated with the gene prediction programs EUGENE [88] and Augustus [89]. The BAC sequences were also searched for genes with BLASTN and BLASTX against the transcripts of SUCEST-FUN database (http://sucest-fun.org/; [15]), the CDS of *S. bicolor, Z. mays* and *O. sativa* from Phytozome v12.0 and the transcripts published by Cardoso-Silva et al. [16]. The BACs were also subjected to BLASTX against Poaceae proteins. The candidate genes were manually annotated using *S. bicolor, O. sativa* and *Z. mays* CDS. The sequences with more than 80% similarity and at least 90% coverage were annotated as genes.

Repetitive content in the BAC clone sequences was identified with the web program LTR_FINDER [90]. Afterward, the BAC sequences were tested by CENSOR [91] against Poaceae.

The phylogenic trees were built by the Neighbor-Joining method [92] with nucleic distances calculated with the Jukes-Cantor model [93] in MEGA 7 software [94]. The Kimura 2-parameter [95] was used as the distance mode.

### Duplication divergence time

The gene contents of *HP600* and *CENP-C* in the duplication regions were compared, and the distance “d” for coding regions was determined by Nei-Gojobori with Jukes-Cantor, available in MEGA 7 software [94]. The divergence times of the sequences shared by the duplicated regions in the BACs were estimated by T = d/2r. The duplicated sequences were used to calculate the pairwise distances (d), and “r” was replaced by the mutation rate of 6.5 × 10-9 mutations per site per year as proposed by Gaut et al. [96]. For the whole duplication, the distance “d” for noncoding regions was determined with the Kimura 2-parameter model and the mutation rate of 1.3 × 10-8 mutations per site per year, as described by Ma and Bennetzen [97].

The insertion ages of the LTR retrotransposons were estimated based on the accumulated number of substitutions between the two LTRs (d) [98], using the mutation rate of 1.3 × 10-8 mutations per site per year, as described by Ma and Bennetzen [97].

### Gene expression

The transcriptomes of the sugarcane variety SP80-3280 from the roots, shoots and stalks were mapped on *HP600* and *CENP-C* (NCBI SRR7274987), and the set of transcripts was used for the transcription analyses. The reads from the sugarcane transcriptomes were mapped to the reference genes with the Bowtie2 software 2.2.5 [99] with default parameters; low-quality reads and unmapped reads were filtered out (SAMtools -b -F 4); bam files were sorted (SAMtools sort); and only mapped reads to the genes were extracted from the bam files (SAMtools fastq) and recorded in a FASTQ format file. A haplotype was considered to be expressed only when the transcript reads were mapped with 100% similarity. SNPs not found in the dataset were searched in the SP80-3280 transcriptomes from Vettore et al. [15], Talbert et al. [33] and Cardoso-Silva et al. [16] to verify the SNP presence in transcripts, but they were not used in the expression analysis.

To test whether the haplotypes had the same proportional ratio in the genome and transcriptome, the transcripts were mapped against one haplotype of the *HP600* haplotypes in Region01 and *CENP-C* with a 90% similarity, and the SNPs found in the transcripts were identified and the coverage and raw variant reads count used to verify the presence of SNPs not found in BACs. An SNP was considered present in the transcripts if it was represented by at least six transcriptome reads [100].

We assumed that one haplotype of each region was missing and tested two genomic frequencies for comparison with the transcriptome sequences: (1) the missing haplotype had a higher frequency of the SNP, and (2) the missing haplotype had a lower frequency of the SNP. When the SNP was not found in the genomic data, we assumed that only the missing haplotype contained the variant SNP.

The frequency of the genomic data was used to test the transcriptome data with R Studio [101] and the exact binomial test (*binom.test* - [102-104]). A p-value >= 0.05 is equivalent to a 95% confidence interval for considering the genomic ratio equal to the transcriptome ratio.

### Chromosome number determination and BAC-FISH

The chromosome number determination was performed as described by Guerra [105] with root tips 0.5–1.5 cm in length, treated with 5 N HCl for 20 min. The slides were stained with Giemsa 2% for 15 min. Chromosome number determination was performed for the varieties SP80-3280, SP81-3250, RB83-5486, RB92-5345, IACSP95-3018 and IACSP93-3046. CMA/DAPI coloration was performed by enzymatic digestion as described by Guerra and Souza [106]. The slides were stained with 10 μg/ml DAPI for 30 min and 10 μg/ml CMA for 1 h. Afterward, the slides were stained with 1:1 glycerol/McIlvaine buffer and visualized.

BAC-FISH was performed using the variety SP803280. For mitotic chromosome preparations, root tips 0.5–1.5 cm in length were collected and treated in the dark with p-dichlorobenzene-saturated solution in the dark at room temperature for 2 h, then fixed in a freshly prepared 3:1 mixture (ethanol: glacial acetic acid) at 4°C for 24 h and stored at −20°C until use. After being washed in water, they were digested with the following enzymes: 2% cellulase (w/v) (Serva, Heidelberg, Baden-Wurtemberg State, Germany), 20% pectinase (v/v) (Sigma, Munich, Baviera State, Germany) and 1% Macerozyme (w/v) (Sigma) at 37°C for 1 h-2 h [107]. The meristems were squashed in a drop of 45% acetic acid and fixed in liquid nitrogen for 15 min. After air-drying, slides with good metaphase chromosome spreads were stored at −20°C.

The BACs Shy064N22 and Shy048L15, both from the BAC library for the SP80-3280 variety, were used as probes. The probes were labeled with digoxigenin-11-dUTP (Roche) by nick translation. Bacterial artificial chromosome-fluorescence in situ hybridization (BAC-FISH) was performed as described by Schwarzacher and Heslop-Harrison [108] with minor modifications. The *C*_o_*t*-100 fraction of the sugarcane variety SP80-3280 genomic DNA, which was used to block repetitive sequences, was prepared according to Zwick et al. [109]. Preparations were counterstained and mounted with 2 µg/ml DAPI in Vectashield (Vector, Burlingame, CA, USA).

The sugarcane metaphase chromosomes were observed and photographed, depending on the procedure, with transmitted light or epifluorescence under an Olympus BX61 microscope equipped with the appropriate filter sets (Olympus, Shinjuku-ku, Tokyo, Japan) and a JAI® CV-M4 + CL monochromatic digital camera (JAI, Barrington, N.J., USA). Digital images were imported into Photoshop 7.0 (Adobe, San Jose, Calif., USA) for pseudocoloration and final processing.

### Genetic map construction

The BAC haplotypes were used to identify 44 sugarcane SNPs in the *HP600* and *CENP-C* exons. The SNP genotyping method was based on MALDI-TOF analysis performed on a mass spectrometer platform from Sequenom Inc.^®^ as described by Garcia et al. [43]. The mapping population consisted of 151 full siblings derived from a cross between the SP80-3280 (female parent) and RB835486 (male parent) sugarcane cultivars, and the genetic map was constructed as described by Balsalobre et al. [21].

## List of abbreviations

ºC: Celsius degrees
BACs: Bacterial Artificial Chromosomes
BES: BAC ends sequencing
Bp: base pairs
BRIX: Soluble solid content
CDS: Coding DNA sequence
CenH3: histone H3
CENP-C: Centromere protein C
CMA: A3 Chromomycin
DAPI: 4’,6-diamidino-2-phenylindole
DNA: Deoxyribonucleic acid
EST: Expressed Sequence Tag
FISH: Fluorescent in situ hybridization
Gb: Giga-base pairs
GBS: Genotyping by sequencing
HMW: High-molecular-weight
Kb: Kilo-base pairs
QTL: Quantitative Trait Loci
LTR: Long Terminal Repeat
Min: Minutes
Mya: Millions Years Ago
PCR: Polymerase chain reaction
PFGE: Pulsed Field Gel Electrophoresis
RNA: Ribonucleic acid
RNAseq: RNA sequencing
Sec: Seconds
SNP: Single Nucleotide Polymorphism
TEs: Transposable Elements

## Declarations

### Ethics approval and consent to participate

Not applicable

### Consent for publication

Not applicable

### Availability of data and materials

Sequence data from this article can be found in the EMBL/GenBank data libraries under the following accession number(s):

BAC sequences: from MH463467 to MH463488.
RNAseq subset data: SRR7274987

### Competing interests

The authors declare that they have no competing interests

### Funding

This study was supported by the São Paulo Research Foundation (FAPESP) (2008/52197-4) and Coordination for the Improvement of Higher Education Personnel (CAPES, Computational Biology Program). The first author was supported by a FAPESP Ph.D. fellowship from FAPESP (2010/50119-6). The third and fourth authors were supported by PD fellowships (MM 2014/11482-9 and CC-S 2015/16399-5). AP received a research fellowship from the National Council for Scientific and Technological Development (CNPq).

## Authors’ contributions

APS, DAS, ERFM, HB, MV, and AAFG designed the study; AB, DAS, HH, JF, MC, MCM, MVRC, ND, NR and SV performed the research; CBC-S, DAS, GSP, MV, M-AVS and RV contributed new analytical/computational tools; AAFG, AB, APS, CBC-S, DAS, GSP, HB, LRP, MCM, MGAL, MS, MV, and SV analyzed the data; and DAS, MV and APS wrote the paper. All authors critically read the text and approved the manuscript.

## Acknowledgments

We gratefully acknowledge Gabriel Rodrigues Alves Margarido for statistical support.

## Supplemental figures and tables Content

**Supplemental Figure 1.** BAC-end locations in the *Sorghum* genome according to BLASTn analysis.

**Supplemental Figure 2.** BAC BLASTn analysis against sugarcane genome contigs. **Supplemental Figure 3.** Schematic representation of phylogenetics and physical duplications.

**Supplemental Figure 4.** Evolutionary relationships of the gene Sobic.008G134700. **Supplemental Figure 5.** Evolutionary relationships of HP600 and CENP-C. **Supplemental Figure 6.** Mitotic metaphases of the sugarcane varieties.

**Supplemental Figure 7.** CENP-C motifs alignments. **Supplemental Table 1.** BAC assembly and annotation. **Supplemental Table 2.** Orthologous genes from Region01. **Supplemental Table 3.** Orthologous genes from Region02.

**Supplemental Table 4.** Number of SNPs found in CENP-C and HP600.

**Supplemental Table 5.** Number of SNPs found in duplicated regions.

**Supplemental Table 6.** Chromosome counts.

**Supplemental Table 7.** Sequenom iPLEX MassARRAY® primers.

